# Personalized alpha-tACS targeting left posterior parietal cortex modulates visuo-spatial attention and posterior evoked EEG activity

**DOI:** 10.1101/2023.01.04.522700

**Authors:** Jan-Ole Radecke, Marina Fiene, Jonas Misselhorn, Christoph S. Herrmann, Andreas K. Engel, Carsten H. Wolters, Till R. Schneider

## Abstract

**Background:** Covert visuo-spatial attention is marked by the anticipatory lateralization of neuronal alpha activity in the posterior parietal cortex. Previous applications of transcranial alternating current stimulation (tACS) at the alpha frequency, however, were inconclusive regarding the causal contribution of oscillatory activity during visuo-spatial attention.

**Objective:** Attentional shifts of behavior and electroencephalography (EEG) after-effects were assessed in a cued visuo-spatial attention paradigm. We hypothesized that parietal alpha-tACS facilitates attention in the ipsilateral visual hemifield. Furthermore, we assumed that modulations of behavior and neurophysiology are related to individual electric field simulations.

**Methods:** We applied personalized tACS at alpha and gamma frequencies to elucidate the role of oscillatory neuronal activity for visuo-spatial attention. Personalized tACS montages were algorithmically optimized to target individual left and right parietal regions that were defined by an EEG localizer.

**Results:** Behavioral performance in the left hemifield was specifically increased by alpha-tACS compared to gamma-tACS targeting the left parietal cortex. This hemisphere-specific effect was observed despite the symmetry of simulated electric fields. In addition, visual event-related potential (ERP) amplitudes showed a reduced lateralization over posterior sites induced by left alpha-tACS. Neuronal sources of this effect were localized in the left premotor cortex. Interestingly, accuracy modulations induced by left parietal alpha-tACS were directly related to electric field magnitudes in the left premotor cortex.

**Conclusion:** Overall, results corroborate the notion that alpha lateralization plays a causal role in covert visuo-spatial attention and indicate an increased susceptibility of parietal and premotor brain regions of the left dorsal attention network to subtle tACS-neuromodulation.

## Introduction

Shifts of covert visuo-spatial attention have been repeatedly associated with a lateralization of neuronal alpha activity along the dorsal attention network [1–3]. Specifically, an increase of cue-related neuronal alpha power has been described in middle and superior occipital cortex, in posterior parietal cortex along the intraparietal sulcus (IPS), as well as premotor regions in the cerebral hemisphere ipsilateral to the attended hemifield, relative to the contralateral hemisphere [1,2]. This activity projects to posterior sensors in magnetoencephalography (MEG) [3–5,see also 6] and electroencephalography (EEG) studies [7–12] and has been related to the active inhibition of unattended space [9–14]. In parallel, cue event-related potentials (ERPs) showed amplitude variations that were increased over posterior sensors ipsilateral to the attended hemifield [6,15,cf. 16]. In contrast, in response to subsequent visual target stimuli, a relative increase of posterior neuronal gamma activity [1,2] and ERP amplitudes [17–19] contralateral to the attended hemifield has been described, reflecting the facilitated processing of attended stimuli [20,21].

To elucidate the role of neuronal alpha oscillations during visuo-spatial attention beyond correlative evidence, transcranial alternating current stimulation (tACS) can be applied to modulate neuronal dynamics, thereby affecting neuronal synchrony and power at the stimulation frequency [22–24]. Especially tACS in the alpha frequency band has been reported to specifically modulate cortical alpha power [23], showing after-effects that outlast the actual stimulation period [25–29]. During visuo-spatial attention experiments, tACS in the alpha frequency range has been repeatedly applied over the left [30–33] or right parietal cortex [31,34–37]. However, the observed behavioral tACS-effects showed limited replicability, hampering the interpretation of neuronal alpha activity as being causal for visuo-spatial attention [32,34,36]. In none of these studies, individual stimulation targets or electric field properties were estimated to validate the potential efficacy of tACS.

In a series of simulations of transcranial electric fields using the finite element method (FEM), interindividual anatomical variability, and thus variability in the magnitude, spatial extent, and orientation of the induced electric field, was identified as a key factor limiting the effects of transcranial electrical stimulation [38–45]. Only recently, the topology and magnitude of individual electric fields have been reported to correlate with the strength of tACS-modulations of neuronal activity [23,46]. Thus, by using algorithmic optimization of individual stimulation montages, personalized tACS has the potential to increase control over the topology and orientation of the electric fields relative to a given stimulation target [47,48]. In addition, this approach allows the post-hoc analysis of the estimated electric fields in conjunction with behavioral or neurophysiological outcome measures of tACS [23,45,cf. 46,48].

Here, we present an application of personalized alpha-tACS, specifically targeting individual sources of neuronal alpha power in the left and right parietal cortices. Parietal alpha power sources were defined based on individual localizer data recorded with high-density EEG. Individual FEM head models were utilized for EEG source imaging, simulations of transcranial electric fields and algorithmic optimization of tACS montages. The posterior parietal cortex along the IPS was chosen as stimulation target as it acts as an important hub within the bilateral dorsal attention network [2,49–51]. Gamma-tACS was applied as a control condition, expecting antagonistic effects compared to alpha-tACS [31,52,53]. In a covert visuo-spatial attention paradigm we investigated tACS modulation of behavior and tACS after-effects in the EEG, as well as their relation to individual electric field simulations.

We hypothesized that the application of personalized alpha-tACS may increase the intrinsic neuronal alpha power within the targeted left or right parietal cortex, thereby facilitating active inhibition of attended stimuli in the visual hemifield contralateral to the targeted hemisphere. This is expected to lead to a relative facilitation of behavior in response to stimuli presented ipsilateral to the hemisphere targeted by alpha-tACS. Based on previous evidence [25–29], we expected that this tACS-modulation may not only be observed during tACS (tACS_ON_), but also elicit after-effects on the behavioral and neurophysiological level (tACS_OFF_).

## Materials and methods

### Participants and procedure

Twenty-two right-handed participants (12 female, 10 male, 27.7 ± 4.2 years [range 20 to 38]) were included in this study. All participants reported no history of neurological or psychiatric disorders and had normal or corrected-to-normal visual acuity and normal hearing. Participants were reimbursed for participation, gave written informed consent in line with the declaration of Helsinki and the protocol was approved by the ethics committee of the Hamburg Medical Association (Ärztekammer Hamburg, PV5338). During four pseudo-randomized sessions, personalized alpha- or gamma-tACS was applied targeting either the left or the right parietal cortex, while participants completed a cued visuo-spatial attention task (Fig. 1). Detailed descriptions of the methods are provided in the supplementary materials.

**Figure 1.**
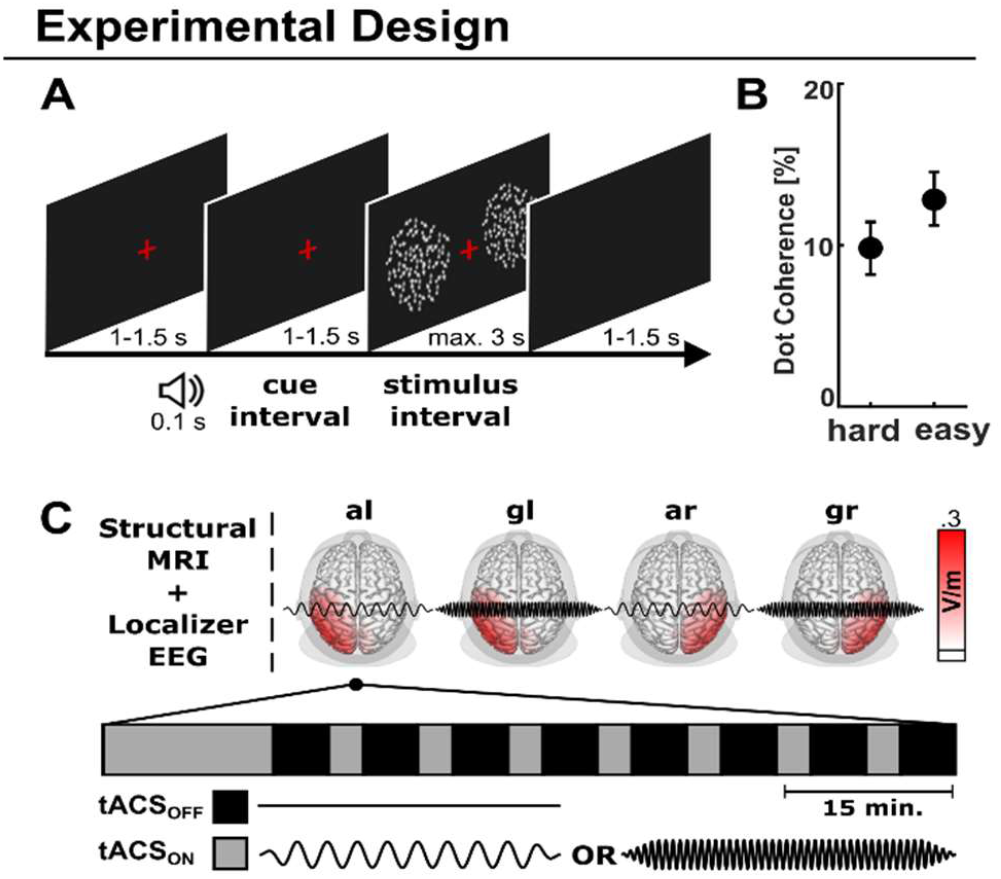
Experimental Design. **A)** A cued visuo-spatial attention paradigm was employed. In each trial, a baseline period was followed by a tone that indicated whether to attend the left or right hemifield during the following cue-stimulus interval. Bilateral random dot kinematograms were presented for up to 3 s, followed by the inter-trial interval. Participants indicated via button press whether the random dots moved up- or downwards in the attended hemifield. **B)** Percentage of dots moving coherently either up- or downwards. Random dots were presented with two difficulties relative to the individually titrated threshold. Mean± standard deviations are depicted. **C)** Top: In a full within subject design, personalized tACS-montages were estimated using structural MRI data and localizer EEG data. Structural MRI data were employed to build realistic headmodels and optimize tACS-montages to target individual alpha sources in the left and right parietal cortex. Four tACS conditions (alpha-left, al; gamma-left, gl; alpha-right, ar; gamma-right, gr; counter-balanced across participants) were applied targeting either the left or right parietal cortex using alpha-tACS (10 Hz) or gamma-tACS (47.1 Hz). Average electric field magnitudes are shown, interpolated on the cortical surface of the MNI brain, viewed from top. Bottom: During each tACS-session, an intermittent stimulation protocol was employed. After an initial 15 min tACS_ON_ interval, short intervals without tACS (tACS_OFF_) were interleaved by short tACS_ON_ intervals (breaks are not shown). During all tACS_ON_ and tACS_OFF_ intervals participants conducted the cued visuo-spatial attention task.

### Cued visuo-spatial attention paradigm

A cued visuo-spatial attention paradigm was utilized to probe participants when attending to the left versus the right hemifield. In each trial, participants were presented with one of two sinusoidal auditory cue stimuli (440 Hz or 880 Hz) [cf. 9]. Cues indicated participants to shift their attention to either the left or the right hemifield while focusing on a red central fixation cross. After a delay period, bilateral random dot kinematograms were presented [1,4,cf. 54,55]. Random dots moved with 11.5°/s with a proportion of dots coherently moving upwards or downwards at individually determined coherence thresholds (Fig. 1A). Participants indicated via button press (The Blackbox Toolkit Ltd., UK) whether the random dots moved up- or downwards in the attended hemifield. Across subjects, individual coherence levels were defined at 9.8 ± 4.2 % (hard) and 12.8 ± 4.2 % (easy; M ± SD; Fig. 1B) using an adaptive procedure [56].

Overall, 400 trials were presented in 8 blocks during the localizer session, while EEG was recorded. During each of the four tACS sessions, 712 trials were presented in 16 blocks, while tACS was applied in an intermittent stimulation protocol (312/712 trials; tACS_ON_) (Fig. 1C). EEG was recorded during non-tACS sequences (400/712 trials; tACS_OFF_). During all sessions, participants were seated inside a dimly-lit electromagnetically shielded booth in front of a computer screen. Custom MATLAB scripts (The Mathworks Ltd., USA) using the Psychophysics Toolbox [57,58] were employed for stimulus presentation.

### EEG data acquisition

EEG data were digitized at a sampling rate of 5 kHz using a BrainAmp EEG amplifier system (BrainProducts, Germany) with an analog filter between 0.016 and 250 Hz and the lab streaming layer (https://labstreaminglayer.org). 126 passive Ag/AgCl electrodes were placed in an equidistant layout (Fig. 2H), with the online-reference placed at the nose tip and a fronto-polar ground electrode (Easycap, Germany). Two electrodes were placed below the eyes to record the electrooculogram (EOG). Electrode impedances were kept below 20 kΩ and individual electrode positions were optically registered (Xensor, ANT Neuro, The Netherlands) for electric field simulations, optimization of tACS montages, and EEG source localization.

**Figure 2.**
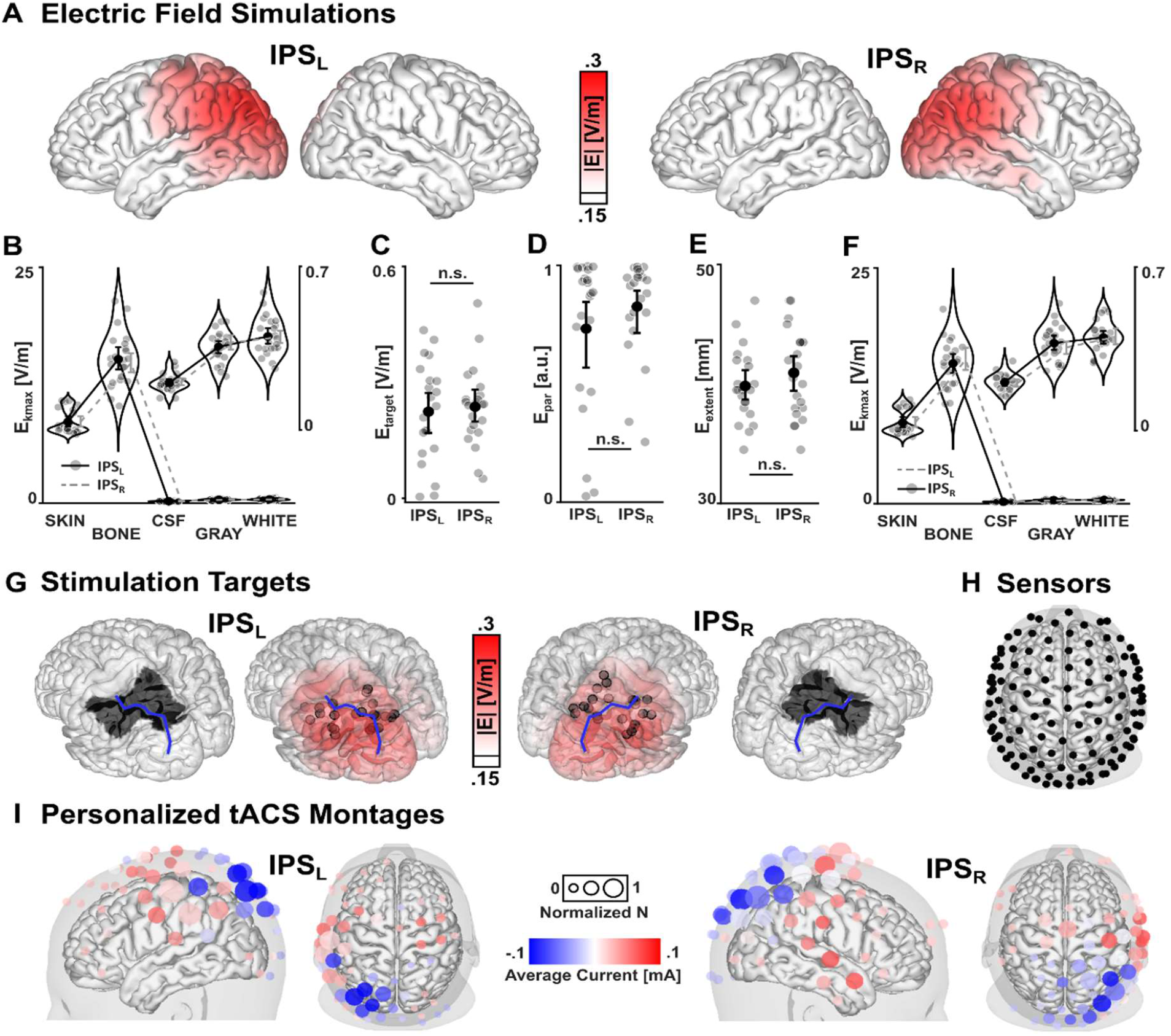
Electric field simulations. **A)** Average magnitude of electric fields targeting the left (IPS_L_) and right (IPS_R_) parietal targets (thresholded at 0.15 V/m). **B)** Unspecific electric field magnitude across tissue type for IPS_L_. Dashed grey lines represent the electric field magnitudes for IPS_R_ for direct comparison. Electric field magnitudes for cerebrospinal fluid (CSF), gray matter (GRAY), and white matter (WHITE) are amplified, due to the scaling differences to SKIN and BONE electric field magnitudes. Electric field magnitudes were similar between IPS_L_ and IPS_R_ across all tissue types. **C)** Target electric field magnitude, **D)** parallelity between electric field orientation and target orientation in the stimulation target, and **E)** spatial extent of electric fields are comparable between IPS_L_ and IPS_R_. **F)** Unspecific electric field magnitude across tissue type for IPS_R_. Dashed grey lines represent the electric field magnitudes for IPS_L_ for direct comparison. Electric field magnitudes were similar between IPS_L_ and IPS_R_ across all tissue types. **G)** Anatomical regions of interest for stimulation target definition (inferior and superior parietal cortex), interpolated on the cortical surface (left and right regions of interest are marked by black patches; left and right intraparietal sulci are marked by blue lines). The inner two plots depict the individual stimulation target coordinates of alpha total power along the intraparietal sulcus (black circles), relative to the average electric field magnitude interpolated on the cortical surface of a standard brain. **H)** Electrode positions from the EEG layout plotted together with the scalp and cortical surface of a standard brain, viewed from the top. The same 126 electrode positions were used for optimization of tACS-montages. **I)** Grand average representation of individual tACS-montages. Circle sizes represent the frequency that each electrode was used for stimulation, normalized to the number of participants. Color-coding represent the average current applied to each electrode. The electrode montage is shown relative to the scalp and cortical surface of a standard brain. n.s. = not significant. Individual values and bootstrapped mean and 95%- confidence intervals are depicted in B) to F).

### MRI data acquisition and FEM head model generation

For each subject structural T1 and T2-weighted magnetic resonance images (MRI) were recorded with a 3T MR-scanner and a 64-channel head coil at an isotropic voxel resolution of 1×1×1 mm (Siemens Magnetom Prisma, Germany). Both, T1 and T2 images were acquired with an MP-RAGE pulse sequence (T1: TR/TE/TI/FA = 2300 ms/ 2.98 ms/ 1100 ms/ 9°, FoV = 192 × 256 × 256 mm; T2: TR/TE = 3200 ms/ 408 ms, FoV = 192 × 256 × 256 mm).

Integrating T1 and T2 imaging data, six compartments were segmented using SPM12-based segmentation (www.fil.ion.ucl.ac.uk/spm) and custom image post-processing including Boolean and morphological operations [44,47,59,60] (see [47] for a detailed description of the procedure). Finally, for each subject isotropic and geometry-adapted hexahedral FEM head models were computed and utilized for the simulation of electric fields induced by tACS, as well as for EEG source localization [42,61]. Individually registered electrode positions from the EEG layout were simulated in the framework of a point electrode model [62].

### Personalized tACS: Preparation and application

Stimulation targets were defined within left and right parietal regions of interest (ROI) at the sites of maximal lateralization of alpha power, based on the EEG localizer data and exact low-resolution electromagnetic tomography (eLORETA) [63] in combination with individual FEM leadfields (Fig. 2G; see Supplement for a details). Target locations and orientations were then used to compute personalized tACS montages.

The Distributed Constrained Maximum Intensity (DCMI) algorithm was utilized [64,65] for the individual targeting of tACS montages, based on the individual 126 electrode positions (Fig. 2H) and the respective six compartment FEM head models with 3.67 ± 0.31 million nodes (see [47] and Supplement for details). In short, the DCMI maximizes the electric field intensity along the orientation of the stimulation target (directionality), while including a parameter that allows to distribute the injected current across stimulation electrodes. In a two-step procedure the number of stimulation electrodes was fixed to six electrodes. The maximal current applied to each electrode was limited to 0.95 mA to reduce potential tactile perception of electrical stimulation.

In four sessions, tACS was applied in an intermittent electrical stimulation protocol either targeting the left or right IPS in the alpha (10 Hz) or gamma frequency (47.1 Hz) [cf. 31,53] (Fig. 1C), resulting in four tACS conditions (alpha-left; gamma-left; alpha-right; gamma-right). A Starstim device (Neuroelectrics, Spain) and Ag/AgCl stimulation electrodes (NG Pistim) with a surface of 3.14 cm^2^ were utilized for stimulation. During each tACS-session, six EEG electrodes from the 126-channel layout (Fig. 2H) were replaced by stimulation electrodes of the personalized tACS-montage (see Supplement). tACS started with 15 min of stimulation (“warmup”), before eight tACS_OFF_ blocks without stimulation (8x 4.5 min) were conducted interleaved with seven short stimulation blocks (7x 3 min, tACS_ON_). This procedure allowed the intermittent recording of EEG data free of electrical tACS-artifacts to analyze stimulation aftereffects during tACS_OFF_ intervals. Gamma-tACS at 47.1 Hz was chosen as a control condition to assess the frequency specificity of tACS effects at a frequency that is not a multiple of 10 Hz [cf. 31,53]. Further, the application of tACS targeting homologue brain areas in the left and right parietal cortex allows the assessment of the spatial specificity of tACS effects [cf. 66,67]. The assessment of the localizer data allows the comparison of the active stimulation conditions to a no-stimulation condition. Since the occurrence of stimulation side-effects is commonly highly variable across participants, tACS was applied with either 1.5 or 2 mA zero-to-peak to minimize the occurrence of phosphenes or transcutaneous side-effects during stimulation. In addition, anesthetic creme (2.5 % lidocaine, 2.5 % prilocaine) was applied to reduce transcutaneous sensations during electrical stimulation [68].

### Analysis of electric field simulations

Personalized electric field simulations were computed targeting either the left (IPS_L_) or the right IPS (IPS_R_). Electric field simulations for alpha- and gamma-tACS were equivalent (quasi-static approximation). To compare electric field simulations between the left and right hemisphere, the electric field magnitude was estimated for each of five tissue types (SKIN, BONE, CSF, GRAY, WHITE) by averaging the 10000 nodes with the highest values (E_kmax_) [23] for electric fields targeting IPS_L_ and IPS_R_, respectively. For each target, we computed the parallelity (E_par_) between the stimulation target orientation vector and the target electric field orientation vector and the target intensity (E_target_) corrected for the parallelity with the stimulation target vector (directionality [64]). Similarly, non-target directionality (E_non-target_) was defined contralateral to the stimulation target. Furthermore, the spatial extent of the electric field relative to the stimulation target (E_extent_) was analyzed [47]. For illustration, individual electric fields were interpolated on a common MNI cortical grid and averaged across subjects for IPS_L_ and IPS_R_, respectively (Fig. 2A; see Supplement for details).

E_kmax_ measures were statistically analyzed in a repeated-measures analysis of variance (ANOVA) including the factors Stimulation Side [IPS_L_, IPS_R_] and Tissue [SKIN, BONE, CSF, GRAY, WHITE]. Target-specific measures (E_target_, E_par_, E_extent_) were tested with paired t-tests to evaluate differences between electric field simulations between IPS_L_ and IPS_R_.

### Behavioral data analysis

Behavioral data were analyzed with respect to performance differences between trials in which participants attended the left (attend_L_) versus the right hemifield (attend_R_). Median reaction times (RTs), as well as sensitivity index *d’* and response bias ln(β) were computed [69], separately for each attention side (attend_L_ and attend_R_), for the tACS_ON_ and tACS_OFF_ intervals, as well as for the first and second half of the experiment. Parameters for the tACS_ON_ intervals were computed for the warmup interval (ON_1_) and integrated across all subsequent tACS_ON_ blocks (ON_2_). For tACS_OFF_ intervals of the first four (OFF_1_) and last four blocks (OFF_2_) were integrated, see Fig. 1C). During the localizer session, only OFF_1_ and OFF_2_ blocks were computed, since no tACS was applied. HITs were defined as probabilities of correct UP responses and false positives (FPs) as probabilities of incorrect DOWN responses (see Supplement for details). The behavioral lateralization during the localizer session was tested with repeated-measures ANOVAs including the factors Block [OFF_1_, OFF_2_] and Attention Side [attendL, attend_R_],separately computed for *d’*, ln(β) and RTs. A similar analysis was conducted to test behavioral lateralization for the four tACS sessions. Therefore, attention contrasts (attend_L_ - attend_R_) were computed for each parameter and stimulation condition. For these contrasts, repeated-measures ANOVAs were computed including the factors Block [ON_1_, ON_2_, OFF_1_, OFF_2_], Stimulation Frequency [alpha, gamma] and Stimulation Side [IPS_L_, IPS_R_], separately for *d’*, ln(β) and RTs.

### EEG data analysis

Due to electrical contamination of the EEG signal during tACS application [70,71], only tACS_OFF_ artifact-free EEG data were analyzed (Fig. 1C) for the four tACS sessions (alpha-left, al; gamma-left, gl; alpha-right, ar; gamma-right, gr). EEG data from the localizer session were analyzed in a similar way to illustrate EEG activity in the absence of tACS during visuo-spatial attention. EEG data were analyzed using MATLAB (The Mathworks Ltd., USA) including the EEGLAB [72], FieldTrip [73] and METH [74] toolboxes, as well as custom scripts.

#### Preprocessing of EEG data

Continuous EEG data were down-sampled to 500 Hz and highpass-filtered at 0.3 Hz half-amplitude cutoff (transition bandwidth = 0.6 Hz). The EEG data were epoched to cue and stimulus onset, respectively (−1 to 1 s), artifactual channels were removed (0.3 ± 1 channels rejected, M ± SD) and EEG epochs holding residual tACS or non-stereotyped artifacts were rejected. A lowpass-filter was applied at 35 Hz to asses low frequency oscillatory brain activity and ERPs (0.3 - 35 Hz). To control for eye movement, bipolar EOG channels were computed for horizontal and vertical eye movement. Independent component analysis (ICA) components related to eye-blinks, electrocardiogram and electrical noise were identified based on topographies, spectra, temporal dynamics, as well as the relation of each component to the EOG [75] and the respective ICA weights were set to zero (11.8 ± 4.4 ICs were rejected, M ± SD). Finally, the data were re-referenced to common average reference and missing channels were interpolated using a spherical spline.

#### EEG spectral analyses

Sensor space alpha total power was computed for the cue interval (−0.75 to 0 s relative to stimulus onset). Power analysis was centered at 10 ± 2 Hz using two Slepian tapers. Results were averaged across electrodes for two posterior electrode clusters of interest in sensor space (left posterior, lp; right posterior, rp; Fig. 3D). eLORETA was utilized to estimate source alpha power in the cue interval (−0.75 to 0 s relative to stimulus onset) along the dominant orientation [76]. A laterality index (LI) was computed as 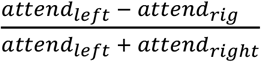 for every grid point. For the localizer, a repeated-measures ANOVA was used for the statistical analysis of sensor-level alpha power including the factors Electrode Cluster [lp, rp] and Attention Side [attend_L_, attend_R_]. Separately, a repeated-measures ANOVA was computed to test for tACS-modulations of alpha power including the factors Stimulation Frequency [alpha, gamma], Stimulation Side [IPS_L_, IPS_R_], Electrode Cluster [lp, rp], and Attention Side [attend_L_, attend_R_]. In case of significant differences on sensor-level, source-level z-scores (uncorrected) were computed, contrasting source estimates of attendL and attend_R_ within each experimental condition (loc, al, gl, ar, gr).

**Figure 3.**
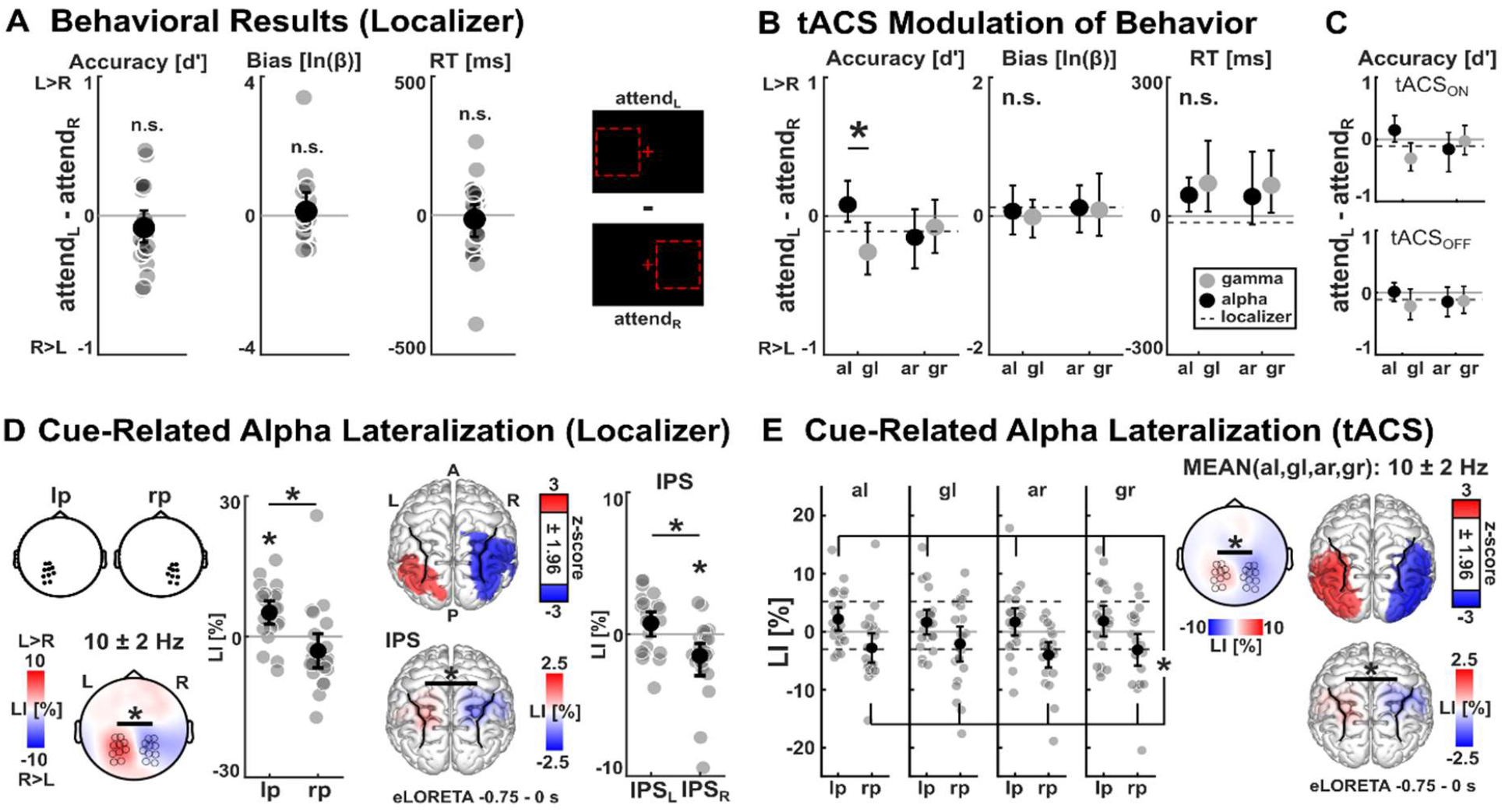
Attentional lateralization of behavior and EEG alpha power. **A)** Left: No differences of accuracies, response bias, or reaction times between attend_L_ and attend_R_ were observed during the localizer session. Individual values (attend_L_-attend_R_) and bootstrapped mean ± 95%-confidence interval are depicted. Right: Illustration of the hypothesized shift of attention in the attend_L_ and attend_R_ conditions during the cue-stimulus interval (see Fig. 1A). **B)** A significant difference was observed between left alpha-tACS (al) and left gamma-tACS (gl) on attend_L_-attend_R_ accuracy differences. No such difference was observed for right alpha-tACS (ar) or right gamma-tACS (gr). Mean values of the localizer session are indicated by dashed black lines for comparisons. **C)** Descriptive accuracy contrasts are shown separately for tACS_ON_ and tACS_OFF_ intervals. **D)** Left: Cue-related alpha total power lateralization contrasting attend_L_ and attend_R_ for the left posterior (lp) and right posterior (rp) electrode cluster and its topographical representation. Positive LI-values (LI = lateralization index) indicate higher alpha power for attend_L_ and negative LI-values indicate higher alpha power for attend_R_. Individual values and bootstrapped mean and 95%- confidence intervals are depicted. Right: Source estimation of the same alpha lateralization (attend_L_ vs. attend_R_) projects to left and right parieto-occipital brain areas along the intraparietal sulcus (z-values thresholded at ± 1.96; positive values indicate higher alpha power for attend_L_). Alpha power lateralization was confirmed for the parietal regions of interest. Individual LI-values and bootstrapped mean ± 95%-confidence interval are depicted. **E)** Cue-related alpha total power, averaged in the left posterior (lp) and right posterior (rp) electrode cluster for the tACS conditions. The alpha lateralization observed during the localizer shown in D) was replicated during the four tACS sessions, yet no tACS-modulation of alpha power lateralization was observed. Mean values of the localizer session are represented by dashed black lines for comparisons. Topographical representations and source estimates averaged across all four tACS-conditions. Individual LI-values and bootstrapped mean ± 95%-confidence interval are depicted, respectively. * represent p < 0.05 corrected for multiple comparisons.

Based on previous literature on visuo-spatial attention [1,2,77], bilateral superior occipital cortices (sOCC), left and right IPS and bilateral middle occipital cortices (mOCC) were defined as posterior ROIs along the dorsal attention network [78]. Power was averaged for all grid points within each region of interest for statistical analysis of source power. To validate alpha lateralization during the localizer on source-level (especially in IPS), a repeated-measures ANOVA was computed. For the localizer, ROI [IPS, mOCC, sOCC], Hemisphere [hemi_L_, hemi_R_], and Attention Side [attend_L_, attend_R_] were defined as factors. Similarly, to assess tACS-modulation effects on alpha power lateralization, a repeated-measures ANOVA was computed including ROI [IPS, mOCC, sOCC], Stimulation Frequency [alpha, gamma], Stimulation Side [IPS_L_, IPS_R_], Hemisphere [hemi_L_, hemi_R_], and Attention Side [attend_L_, attend_R_] as factors.

#### ERP analyses

In addition, visual ERPs were assessed as an indicator of attention-modulated neuronal activity. Sensor-level ERPs were computed in response to random dot stimuli (−0.2 to 0.75 s, relative to stimulus onset), separately for attending to the left and right hemifield, as well as for each stimulation condition. Epochs were averaged and baseline-corrected (−0.2 to 0 s). Difference ERPs were computed by subtracting ERPs of attend_R_ from ERPs of attend_L_ (attend - attend_R_). eLORETA [63] was utilized for source localization of ERPs. eLORETA solutions were computed for the ERP activity, averaged across the respective time window of interest for the ERPs of the localizer and each stimulation condition and attention side. LI was computed as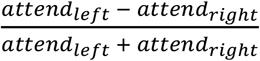 for every grid point based on the source estimates of attend_L_ and attend_R_.

A non-parametric cluster permutation test [79] was conducted to test for differences between ERPs related to attend_L_ and attend_R_ during the stimulus interval of the localizer. The permutation test for the stimulus-related ERPs was applied for the time-window 0 to 0.6 s relative to stimulus-onset and all 126 EEG sensors (paired *t*-tests, 1000 permutations, α_cluster_ = 0.05, α = 0.05, two-sided). In case of significant results for the localizer ERPs, average mean amplitudes of sensor ERPs (attend_L_ and attend_R_) across time-points and electrodes of each cluster (cluster sensors with < 50 time samples and cluster time samples including < 10 sensors were neglected) were submitted to a paired *t*-test to validate subsequent mean amplitude extraction and parametric testing for tACS conditions. These spatiotemporal clusters were then used to assess ERP differences across the four tACS-sessions.

Mean amplitudes of sensor ERPs (attend_L_ and attend_R_) of all stimulation conditions (al, gl, ar, gr) were extracted, averaged over time-points and electrodes of each cluster that was defined by the cluster permutation tests from the localizer session. Mean amplitudes were then conveyed to a repeated-measures ANOVA, including factors Stimulation Frequency [alpha, gamma], Stimulation Side [IPS_L_, IPS_R_], Spatio-Temporal Cluster [left negative, right positive] and Attention Side [attend_L_, attend_R_]. In case of significant differences on sensor-level, source-level *z*-scores (uncorrected) were computed, contrasting sources of attend_L_ and attend_R_ within each experimental condition (loc, al, gl, ar, gr), as well as attend_L_-attend_R_ differences between experimental conditions.

### Correlation analysis

Correlations between behavioral modulations and the simulated transcranial electric field magnitudes were computed to further explore the role of interindividual differences in the applied electric fields. Individual electric field magnitudes were interpolated to a common 5 mm source grid and correlated to the tACS-modulation of behavior (attend_L_-attend_R_ *d’*-contrast), separately for online effects (tACS_ON_) and after-effects (tACS_OFF_). Non-parametric cluster permutation tests [79] were conducted to test for significant Spearman correlations using 1000 permutations (α_cluster_ = 0.01, α = 0.05, two-sided).

For all statistical anlyses IBM SPSS Statistics (IBM Corp., USA) and MATLAB (The Mathworks Ltd., USA; including FieldTrip) were utilized for statistical analyses. Significance levels were set to α = .05. For ANOVAs, Greenhouse-Geisser correction was applied in case the sphericity assumption was violated and follow-up paired *t*-tests or Wilcoxon signed-rank tests (in case of violated normality assumption) were computed for the highest-order interaction or main effects, respectively. Results from *t*-tests were corrected for multiple comparisons using the Bonferroni-Holm correction [80]. In case of significant results, test-values, corrected *p*-values, as well as effect sizes are reported. For cluster permutation tests, cluster *p*-value (corrected) and the number of spatio-temporal samples in the cluster (*n*_clustersize_) are reported for significant effects.

## Results

### Electric field simulations targeting the left and right hemisphere show no difference

Electric field simulations revealed overall cortical electric field magnitudes of E_kmax_ = 0.37 ± 0.06 V/m (GRAY, M ± SD) with highest values in posterior brain regions along the left and right IPS, respectively (Fig. 2A). On average, a reasonable and specific electric field magnitude, anti-/parallel to the stimulation target orientation was observed for IPS_L_ (E_target_ = 0.22 ± 0.03 V/m, E_non-target_ = 0.07 ± 0.02 V/m) and IPS_R_ (E_target_ = 0.24 ± 0.02 V/m, E_non-target_ = 0.06 ± 0.01 V/m, M ± SEM), respectively. The repeated-measures ANOVA of unspecific electric field magnitudes (E_kmax_) across Tissue and Stimulation Side showed a significant main effect of Tissue (*F*_1.2,24.7_ = 733.23, *p* < .0001, 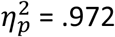), whereas no main or interaction effect including Stimulation Side was observed (all p > .151). Paired t-tests confirmed differences in E_kmax_ between tissues (BONE > SKIN > WHITE/GRAY > CSF, all *t*_21_ >|17.56|, all *p* < 0.001, all *d* > 3.75, except WHITE versus GRAY; Fig. 2B and F, see Supplement). No significant differences were observed between IPS_L_ and IPS_R_ for neither E_target_ (*p* = .645; Fig. 2C), E_par_ (*p* = .186; Fig. 2D), nor E_extent_ (*p* = .237; Fig. 2E). Overall, these results indicate that no differences were observed between the applied tACS electric fields targeting IPS_L_ and IPS_R_. Anatomical target regions and the pooled stimulation target coordinate vectors relative to the average cortical electric field, as well as the stimulation montages are depicted for IPS_L_ and IPS_R_ in Fig. 2 (Fig. 2G and 2I; see Supplement).

### Left alpha-tACS enhances behavioral performance when attending the left hemifield

On average, during the localizer, participants showed hit-rates of 70 ± 9 % and reaction times of 1377 ± 111 ms (M ± SD). As intended, no attentional lateralization was observed, neither of accuracies, response bias, nor reaction times (all interactions and main effects: *p* > .141, Fig. 3A).

During the four tACS sessions, participants showed average hit-rates of 76 ± 2 % (al, M ± SD), 77 ± 2 % (gl), 78 ± 2 % (ar), and 76 ± 2 % (gr), as well as average reaction times of 1408 ± 127 ms (al), 1379 ± 103 ms (gl), 1332 ± 97 ms (ar), and 1382 ± 102 ms (gr). The repeated-measures ANOVA of attend_L_ - attend_R_ *d’*-contrasts (*d’*_al_, *d’*_gl_, *d’*_ar_, *d’*_gr_) revealed a significant interaction of Stimulation Frequency and Stimulation Side (*F*_1,21_ = 9.51, *p* = .006, 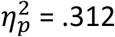), as well as a main effect of Stimulation Frequency (*F*_1,21_ = 4.44, *p* = .047, 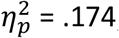; all other main or interaction effects: *p* > .074). Paired t-tests confirmed a significant difference between left alpha-tACS and left gamma-tACS (contrast *d’*_al_ > *d’*_gl_: *t*_21_ = 4.26, *p* = .0014, *d* = .909; Fig. 3B), indicating relatively higher accuracies for left alpha-tACS, when attending to the left hemifield, compared to the right hemifield (al: attend_L_ *d’* = 1.79 ± 0.16, attend_R_ *d’* = 1.72 ± 0.16 ; M ± SEM) and vice versa for left gamma-tACS (gl: attend_L_ *d’* = 1.68 ± 0.18, attend_R_ *d’* = 1.88 ± 0.19). No significant differences were observed comparing *d’* values for any other combination of stimulation conditions (all *p* > .135). Importantly, the non-significant contribution of the factor Block indicates that the behavioral effect observed during tACS_ON_ also translated to tACS_OFF_ intervals, although the difference between al and gl decreased descriptively during tACS_OFF_ (Fig. 3C). Apart from tACS effects on *d’*-contrasts, no significant effects were observed for response bias (all main effects and interactions: *p* > .066). For reaction times, a significant Stimulation Frequency * Block interaction (*F*_2.5,49.2_ = 3.37, *p* = .034, 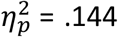; all other main effects and interactions: *p* > .098) was observed. However, follow-up t-tests of reaction times averaged across stimulation frequencies did not reveal significant differences (all *p* > .37).

### No tACS-modulation of cue-related alpha lateralization in EEG after-effects

Sensor-level analysis of cue-related alpha total power during the localizer revealed a significant interaction of Electrode Cluster and Attention Side (*F*_1,21_ =7.38, *p* = .013, 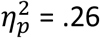), as well as a main effect of Electrode Cluster (*F*_1,21_ = 9.38, *p* = .006, 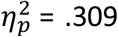), but no main effect of Attention Side (*p* > .356). Paired *t*-tests revealed a significant alpha power difference between attend_L_ and attend_R_ in the left (*t*_21_ = 2.91, *p* = .017, *d* = .62), but not the right posterior electrode cluster (*p* = .128). In addition, a significantly different LI (contrasting attend_L_ and attend_R_) was observed between the two electrode clusters (*t*_21_ = 3.83, *p* = .001, *d* = .816; Fig. 3D, left), indicating enhanced alpha power in left posterior electrodes during attend_L_, compared to attend_R_ and the opposite pattern in right posterior electrodes.

The sources of lateralized alpha power during the localizer cue interval span along the ventral IPS in the left hemisphere and the ventral and posterior IPS in the right hemisphere (Fig. 3D, right). The repeated-measures ANOVA, probing a lateralization of cue-related alpha power on source-level, revealed a ROI * Hemisphere * Attention Side interaction (*F*_1.5,32.1_ = 6.73, *p* = .007, 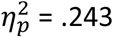), and a Hemisphere * Attention Side interaction (*F*_1,21_ = 9.5, *p* = .006, 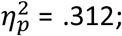; all other main or interaction effect: *p* > .06). Post-hoc Wilcoxon signed-rank tests revealed significant differences between alpha source power between attend_L_ and attend_R_ in right IPS, right sOCC, and bilateral mOCC (Fig. 3D; see Supplement). LIs were different between hemispheres for all three ROIs (all *t*_21_ > 3.61, all *p* < .002, all *d* > .77) and no differences were observed between ROIs within each hemisphere (all *p* > .064).For the tACS conditions, no modulation of cue-related alpha power was observed, neither on sensor-level nor on source-level. However, the repeated-measures ANOVA reproduced the significant interaction of Electrode Cluster and Attention Side (*F*_1,21_ =9.45, *p* = .006, 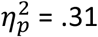), as well as a main effect of Electrode Cluster (*F*_1,21_ = 7.48, *p* = .012, 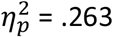) that was already observed during the localizer. No specific tACS-effect was observed (all other main or interaction effects: *p* > .052). Paired *t*-tests confirmed a significant alpha power difference between attend_L_ and attend_R_ (averaged across all stimulation conditions) in the left electrode cluster (*t*_21_ = 2.55, *p* = .019, *d* = .543), but also revealed significant power differences in the right posterior electrode cluster (*t*_21_ = -2.94, *p* = .016, *d* = .626). The LI was significantly different between the two electrode clusters (*t*_21_ = 3.7, *p* = .001, *d* = .789), indicating a relatively increased alpha power in left posterior electrodes when attend_L_ was compared to attend_R_ and the opposite pattern in right posterior electrodes (Fig. 3E).

Averaged across all four stimulation conditions (al, gl, ar, gr), the sources of lateralized alpha power during the cue interval extended from ventral IPS to posterior IPS in the left hemisphere relative to the localizer. Source power was localized to ventral, as well as posterior IPS in the right hemisphere, as illustrated by source-level *z*-scores (Fig. 3E). The repeated-measures ANOVA, probing tACS-modulation of cue-related alpha lateralization on source-level, revealed a ROI * Hemisphere * Attention Side interaction (*F*_1.2,25.5_ = 4.15, *p* = .045, 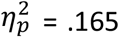), and a Hemisphere * Attention Side interaction (*F*_1,21_ = 12.41, *p* = .002,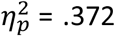), a Stimulation Side * Hemisphere interaction (*F*_1,21_ = 5.3, *p* = .032, 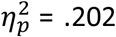), as well as main effects of Stimulation Side (*F*_1,21_ = 4.39, *p* = .048, 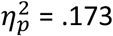) and ROI (*F*_1.5,31.5_ = 3.7, *p* = .048, 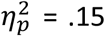; all other main or interaction effect: *p* > .079). Post-hoc paired Wilcoxon signed-rank tests revealed significant differences between alpha source power between attend_L_ and attend_R_ (averaged across all stimulation conditions) in bilateral IPS, bilateral sOCC, and bilateral mOCC (Fig. 3E, see Supplement). LIs were different between hemispheres for all three ROIs (all *t*_21_ > 6.2, all *p* < .001, all *d* > 1.321). Left mOCC showed an increased LI, compared to left IPS (*t*_21_ = -3.2, *p* = .026, *d* = .681) and the right mOCC showed a stronger lateralization, compared to right sOCC (*t*_21_ = 2.87, *p* = .046, *d* = .612).

### Left alpha tACS modulates visual ERP activity in left premotor cortex

During the localizer, attention-related amplitude modulations (attend_L_, attend_R_) were observed in bilateral posterior electrode clusters (lp, rp) for the visual ERPs (Fig. 4A). Visual ERPs strongly varied between attention conditions with more positive amplitudes for attended stimuli in the hemifield contralateral to the respective electrode cluster. Comparing attend_L_ with attend_R_, we observed a significant positive effect (*p* = .001) in a right posterior electrode cluster (*n*_clustersize_ = 4839, 126 to 498 ms; Fig. 4A, top) and a significant negative effect (*p* = .001) in a left centro-posterior electrode cluster (*n*_clustersize_ = 4418, 106 to 450 ms; Fig. 4A, bottom) revealed by cluster permutation tests. Thus, stimulus ERPs were increased in amplitude over the hemisphere contralateral to the attended hemifield.

**Figure 4.**
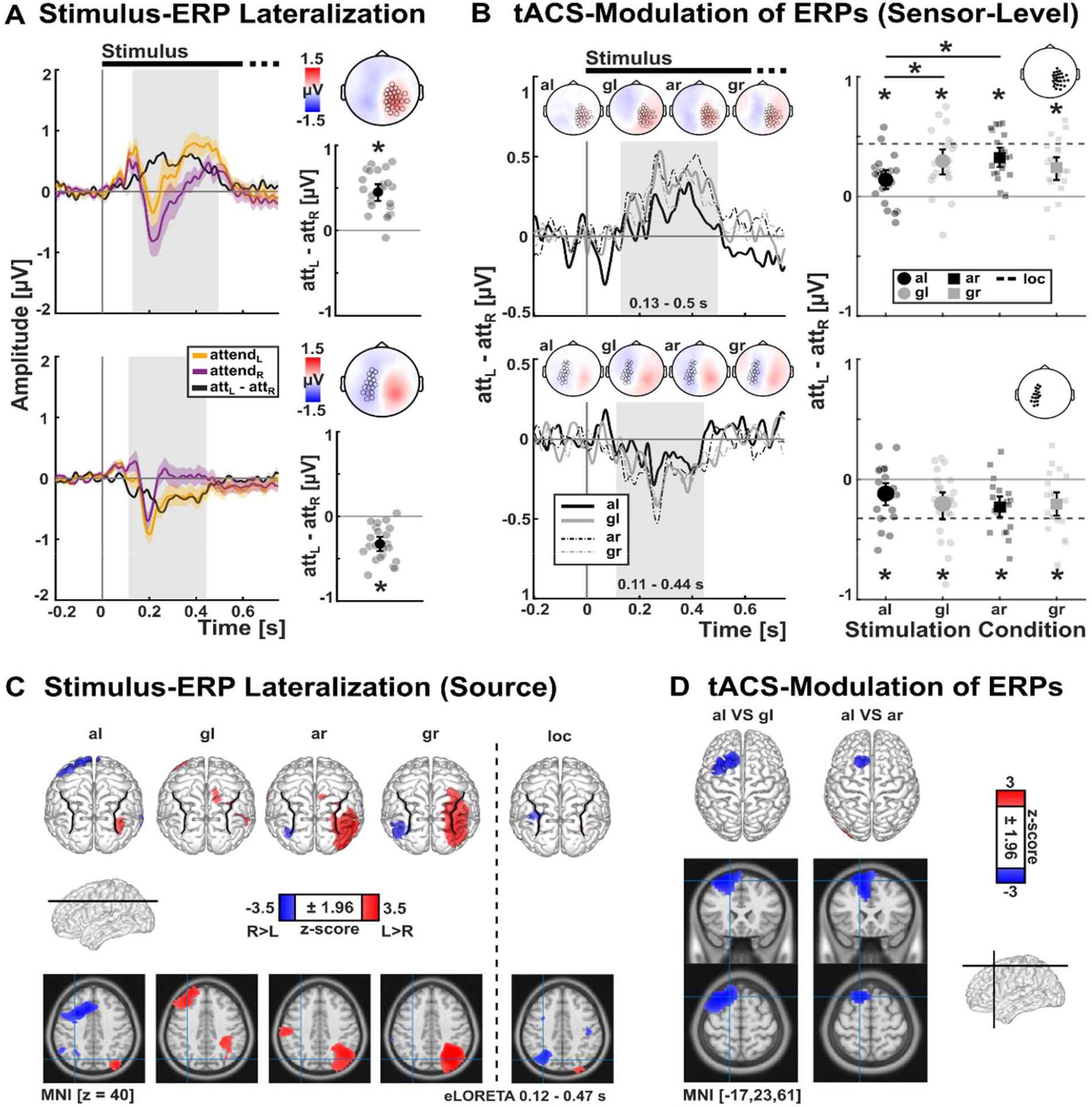
Stimulus-ERP lateralization is modulated by tACS. **A)** During the localizer session, significant stimulus-related amplitude differences between attend_L_ and attend_R_ were identified in a right posterior (top) and a left central-posterior electrode cluster (bottom). Visual ERPs (shading indicates mean ± standard error of the mean), difference ERPs, topographical representations and mean as well as individual ERP amplitudes for the two clusters are presented. Individual amplitude values (grey dots) and bootstrapped mean ± 95%-confidence intervals (black dot and error bars) are depicted. **B)** Left: Difference ERPs (attend_L_-attend_R_) are shown for the four tACS-conditions (alpha-left, al; gamma-left, gl; alpha-right, ar; gamma-right, gr) for the two electrode clusters that were defined during the localizer shown in A). Right: Difference ERP-amplitudes for the right posterior cluster revealed a significant difference between al and gl, as well as al and ar, indicating a relatively reduced ERP lateralization by left alpha tACS. In addition, for all tACS conditions and both clusters, the ERP amplitude differences between attend_L_ vs. attend_R_ were statistically significant. Individual values (attend_L_-attend_R_) and bootstrapped mean ± 95%- confidence intervals are depicted. **C)** Source representations of attend_L_-attend_R_ difference ERPs for all four tACS conditions and the localizer. **D)** Source representations of the significant contrasts between difference ERPs shown in B) show ERP difference contrasts in left premotor cortex when comparing al with gl, as well as al and ar. * represent p < 0.05, corrected for multiple comparisons.

Sensor-level ERPs of all tACS sessions (al, gl, ar, gr) for attend_L_ and attend_R_ conditions were analyzed in the left and right spatio-temporal clusters defined by cluster permutation statistics of the localizer (Fig. 4A). Statistical analysis of stimulus ERPs revealed a significant interaction effect of Stimulation Frequency, Stimulation Side, Spatio-Temporal Cluster and Attention Side (*F*_1,21_ = 10 *p* = .005, 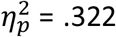), an interaction of Spatio-Temporal Cluster with Attention Side (*F*_1,21_ = 52.44 *p* < .001, 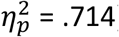) and a main effect of Spatio-Temporal Cluster (*F*_1,21_ = 8.41, *p* < .009, 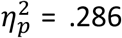; all other *p* > .09). Post-hoc *t*-tests confirmed significant differences between attend_L_ and attend_R_ for all stimulation conditions in both spatio-temporal clusters (all |*t*_21_| > 2.43, all *p* > .024, all *d* > .518; Fig. 4), indicating increased amplitudes in response to stimuli in the contralateral hemifield for all stimulation conditions (Fig. 4B). Descriptively, the difference ERPs spanned the whole latency range of visual P1, N1, P2 and P3 ERP components (see Supplement), peaking between 250-400 ms after stimulus-onset (Fig. 4B).To assess tACS-effects on sensor-level ERPs, follow-up paired *t*-tests were conducted separately for the left and right hemisphere clusters to directly compare attend_L_ - attend_R_ difference ERPs between stimulation conditions. Interestingly, in the right posterior cluster significant differences were revealed between al and gl (*t*_21_ = -2.86, *p* = .047, *d* = .609), as well as between al and ar (*t*_21_ = -5.16, *p* = .0002, *d* = 1.1; Fig. 4B, top). No differences were observed for the other comparisons of difference ERPs between tACS conditions in the right cluster (all *p* > .357), or for any comparison in the left cluster (all *p* > .237; Fig. 4B, bottom).

Sources of ERPs were estimated for attend_L_-attend_R_ differences across stimulation conditions (0.12 to 0.47s relative to stimulus onset), projecting to left and right posterior cortices in all conditions, and to left frontal cortex in conditions al and gl (Fig. 4C). The sources of the ERP differences between al and gl were estimated in the left premotor cortex, specifically extending from left dorsolateral cortex and medial parts of the superior frontal cortex to posterior parts of the middle frontal gyrus and left supplementary motor area (Fig. 4D). Sources of the difference between al and ar were estimated in left premotor cortex, as well as left middle and inferior occipital cortex, including posterior parts of the middle temporal gyrus (Fig. 4D, cf. Fig. 4C).

### Electric field magnitude in left premotor cortex correlates with behavior during left alpha-tACS

In this study, tACS targeting the left parietal cortex (IPS_L_) yielded significant differences of behavioral accuracies between left alpha and left gamma stimulation. In addition, tACS was shown to affect stimulus-evoked neuronal activity in left premotor cortex. Importantly, based on these findings, electric field magnitudes in a cluster in left premotor cortex and adjacent regions were shown to be negatively correlated with behavioral *d’* contrasts after left alpha-tACS (*p* = .001, *n*_clustersize_ = 2695; Fig. 5). Accordingly, if the electric field during left alpha-tACS was higher in left premotor cortex, participants show relatively decreased accuracies for stimuli attended in the left hemifield (i.e., an attention shift to the right hemifield). No significant correlations were observed between the electric field and *d’* contrasts estimated during tACS_ON_, or *d’* contrasts in the left gamma-tACS condition.

**Figure 5.**
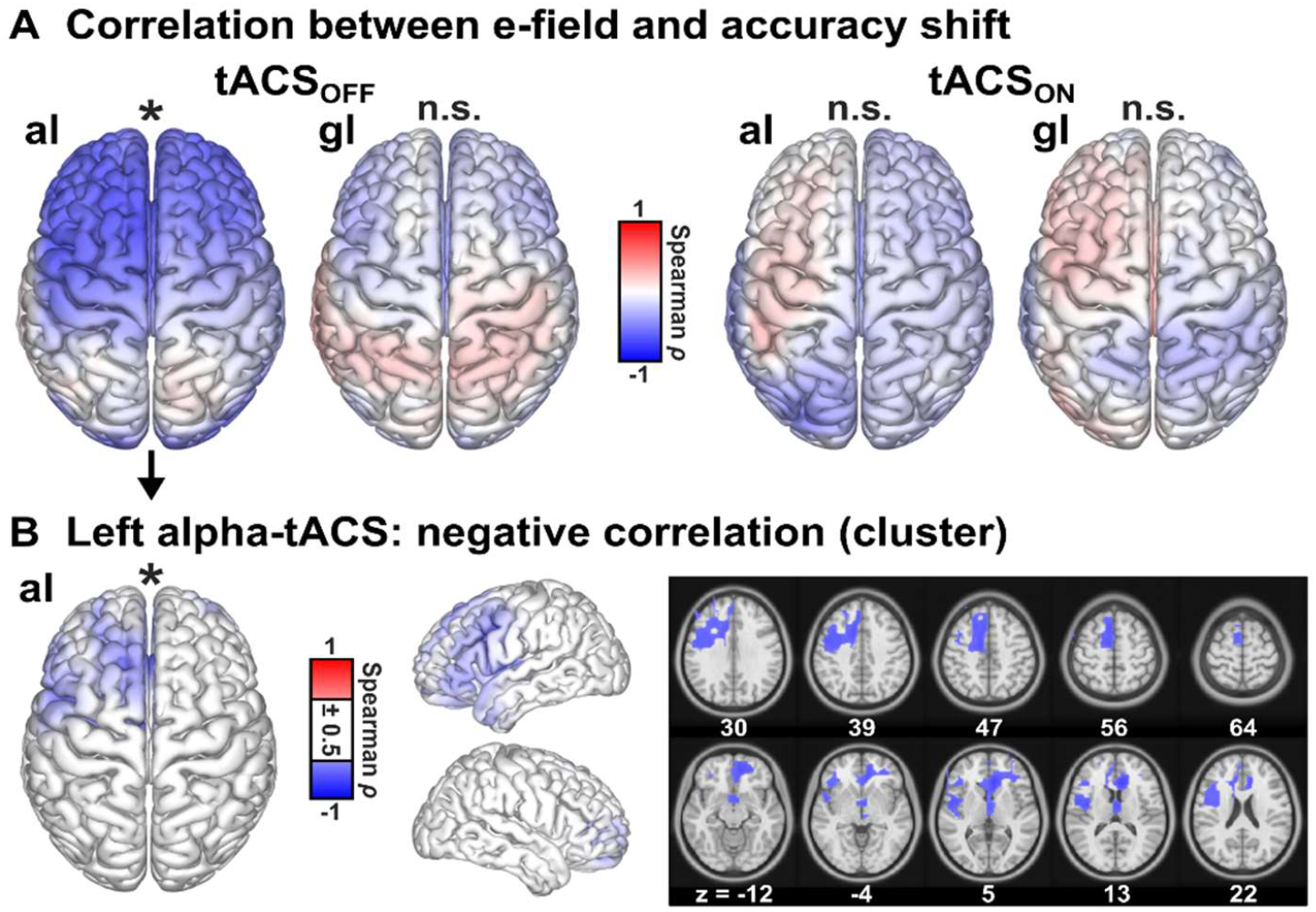
Correlations between accuracies and electric field magnitudes. **A)** Spearman correlations were computed between the behavioral *d’* contrasts (attend_L_-attend_R_) and the whole-brain representation of individual electric fields that target the left parietal cortex (IPS_L_), separately for the left alpha-tACS (al) and left gamma-tACS (gl) conditions and separately for tACS_OFF_ (left) and tACS_ON_ intervals (right). Spearman *ρ*values are shown, interpolated on the cortical surface of the MNI brain. Cluster permutation statistics revealed a negative correlation only for the left alpha-tACS condition during tACS_OFF_ intervals (most left). **B)** A significant negative correlation between the electric field magnitude and the *d’* contrast during tACS_OFF_ was induced by left parietal alpha-tACS, based on a cluster in left premotor cortex. Spearman *ρ*values within the cluster are interpolated on the cortical surface of the MNI brain (left) and in horizontal slices (right). For illustrative reasons, absolute *ρ*values below 0.5 are not shown. Two major foci of the cluster can be identified in left dorsomedial premotor cortex (supplementary motor area) and in left lateral premotor cortex. * represent *p* < 0.05, corrected for multiple comparisons.

## Discussion

Personalized alpha-tACS and gamma-tACS were applied to the left and right posterior parietal cortex during a visuo-spatial attention paradigm using an intermittent stimulation protocol. This procedure allowed the assessment of behavioral tACS modulations, individual electric field simulations, as well as tACS after-effects in the EEG. We showed that personalized alpha-tACS targeted to the left parietal cortex increased accuracies when participants attended the left hemifield relative to the right hemifield, when compared to left gamma-tACS. This behavioral effect was accompanied by a significantly reduced ERP amplitude lateralization in right posterior sensors during left parietal alpha-tACS, compared to left parietal gamma-tACS and right parietal alpha-tACS. EEG source reconstruction located this ERP effect in left premotor cortex. Interestingly, the attentional shift induced by left parietal alpha tACS was dependent on electric field magnitudes in the left premotor cortex.

### Left parietal alpha-versus gamma-tACS induces an attentional shift to the left hemifield

Assuming that neuronal alpha power in the posterior parietal cortex [1,2] can be modulated by tACS, our behavioral finding of a discrimination performance shift to the left hemifield by left alpha tACS compared to left gamma-tACS (Fig. 3B) is in line with previous studies showing that alpha-tACS over the left parieto-occipital cortex facilitates attentional shifts to the ipsilateral hemifield during covert visuo-spatial attention [30,31,33]. Specifically, during covert attention, alpha-tACS over the left parieto-occipital cortex induced faster reaction times in simple discrimination tasks, when attending the left hemifield, relative to the right hemifield [30,31]. No tACS-modulation of RTs was observed during exogenous attention [30,31], or with tACS over right parieto-occipital cortex [31]. Interestingly, the observed shift of accuracies (d’) in our data indicates that neuronal alpha activity can not only be associated with the disengagement and re-allocation of attention in invalidly cued trials [31], but also affects the local perceptual processing in the attended hemifield for valid trials. Moreover, the observed dichotomy of alpha versus gamma tACS in our study has been described previously during visuo-spatial [31] and auditory-spatial attention [53] and can be related to antagonistic effects of neuronal activity in the alpha- and gamma-band [52,81–86].

Here, we substantiate previous findings of non-personalized tACS over parietal cortex by evaluating individual tACS-induced electric fields that explicitly target the left and right parietal cortices. Importantly, the central finding that behavioral tACS-modulations could only be observed after left, but not right, alpha-versus gamma-tACS cannot be explained by differences in applied electric fields (Fig. 3B). Electric fields targeting the left or right parietal cortex were comparable with respect to magnitudes across tissues (Fig. 2A, B and F) and in the stimulation targets (Fig. 2C), the parallelity between the electric field orientations and the stimulation target orientations (Fig. 2D), and the spatial extent of electric fields (Fig. 2E). Interestingly, in a recent MEG-neurofeedback study specifically focusing on the endogenous modulation of visuo-spatial attention, data showed that attention-related alpha lateralization was primarily driven by a modulation of left rather than right posterior alpha activity [87]. This finding was supported by tACS applications that showed specific modulation of endogenous visuo-spatial attention by posterior alpha-tACS over left [30,31,33], but not right hemisphere [31]. Although some studies reported a shift of attention to the right hemifield by tACS over the right parietal cortex [34,35], these results showed limited replicability [34,36,37]. Taken together, our presented data might indicate an increased susceptibility of the left dorsal attention network to subtle tACS-induced neuromodulation during visuo-spatial attention.

### No evidence for outlasting tACS-modulations of cue-related alpha power

During both the localizer experiment and across all four tACS sessions, we observed a pronounced lateralization of alpha oscillatory activity (Fig. 3D and 3E), substantiating previous studies that showed a relative increase of alpha power ipsilateral to the attended hemifield [4,5,7–12] along the intraparietal sulcus [1–3,50] (Fig. 3D and 3E). However, we did not observe the hypothesized modulation of posterior alpha power after-effects by the application of personalized alpha-tACS targeting the left and right parietal cortex, neither at sensor-level (Fig. 3E), nor source-level (Fig. 3E, see Supplement). It is important to note that the analysis of concurrent electrophysiological effects was precluded by strong electrical artifacts during tACS. Therefore, data analysis relied on outlasting effects of stimulation in the tACS_OFF_ intervals. However, after-effects of tACS are associated to lasting neuroplastic changes [25,28] and may differ from entrainment-related online effects [88] that decay quickly after the end of stimulation [89,90]. Thus, although alpha power after-effects were not observed in the present study, this does not preclude an effective online entrainment of alpha rhythms that can have given rise to behavioral modulations. In support of this assumption the behavioral effects were descriptively reduced for tACS_OFF_ compared to tACS_ON_ intervals (Fig. 3B) and may suggest a limited transfer of online tACS-modulation of neuronal alpha power to offline intervals.

### Left alpha-tACS modulates ERP-amplitude lateralization in left premotor cortex

During the assessment of stimulus ERPs, a lateralization of amplitudes was revealed in left and right posterior electrodes that was modulated by left alpha-tACS (Fig. 4). Specifically, the difference stimulus ERPs showed a posterior positivity with a left posterior inversion and a clear peak in the latency range of the P3b ERP component [91], indicating larger amplitudes in the posterior electrodes over the hemisphere contralateral to the attended hemifield (Fig. 4A and 4B). An increase in the posterior P3b amplitude has been proposed to reflect the allocation of top-down attentional resources towards relevant stimuli [92–94], thereby facilitating behavior. In line with our results, visuo-spatial ERP components have been repeatedly shown to increase over posterior scalp regions contralateral to the attended hemifield, indicating the facilitated processing of attended stimuli [17–19]. Critically however, in the present study, the difference ERP amplitudes were reduced during left alpha-tACS (Fig. 4B), while an increased lateralization of accuracies to the left hemifield was observed during the same condition (Fig. 3B). Thus, the ERP amplitudes during left alpha-tACS do not seem to indicate an additional allocation of attentional resources related to the P3b, since that would have been marked by an increased amplitude lateralization. Importantly, P3b sources would be expected in posterior brain regions [94]. In contrast, in our study, eLORETA sources of the ERP amplitude variations and the difference between left alpha- and gamma-tACS were estimated in left premotor cortex for the left alpha-tACS condition (Fig. 4C-D), covering a similar area as described in previous fMRI experiments on visuo-spatial attention [3,95,96]. ERP amplitudes in premotor cortex were relatively decreased when attending to the (ipsilateral) left hemifield during left parietal alpha-tACS (Fig. 4C). The observed ERP modulation in premotor cortex includes the supplementary motor area, which is associated with the preparation of self-initiated movements [97,98] and, more importantly, the preparation of eye movements towards a cued location [96,99,100], tightly linking networks of visuo-spatial attention to oculomotor function [96,101–104]. Furthermore, premotor cortex has been proposed to be tightly coupled with parietal and occipital brain regions during visuo-spatial attention [3,105,106]. In the present study, the attentional shift to the left hemifield induced by left alpha-tACS was accompanied by decreased stimulus ERP amplitudes in the left premotor cortex when attending the left hemifield relative to the right hemifield (left gamma-tACS induced the opposite effects; Fig. 3B and 4C). A similar shift of attention towards the left hemifield has been described when the left premotor cortex was inhibited by repetitive transcranial magnetic stimulation [3]. Since, in the present study, posterior parietal cortex was specifically targeted by tACS, we assume that left alpha-tACS versus gamma-tACS could have modulated the left fronto-parietal network and, thus, stimulus ERP amplitudes in frontal areas. Specifically, our results indicate that tACS might have affected parietal control over premotor areas [cf. 51] or connectivity in the fronto-parietal network [3,35,36,50,107] which gave rise to behavioral attention effects.

### Electric field magnitudes in left premotor cortex are related to behavioral lateralization

Interestingly, we observed a correlation between the electric field magnitude in the left premotor cortex (showing an ERP amplitude modulation by tACS) and the behavioral shift of attention (indexed by *d’*) during the tACS_OFF_ interval after left parietal alpha-tACS (Fig. 5). These results indicate a potential co-stimulation of left premotor cortex when targeting the IPS_L_. Specifically, higher electric field magnitudes in the left premotor cortex were associated with a relative facilitation of accuracies (*d’*) in the right hemifield. Thus, this co-stimulation of left premotor cortex counteracted the attentional shift to the left hemifield. These results indicate that the co-stimulation of left premotor and left parietal cortex affected the connectivity in the fronto-parietal network [3,35,36,50,107] differently compared to the predominantly parietal stimulation.

Co-stimulation of brain regions apart from the tACS target region are inevitable when optimizing electric fields with regard to target intensity. As electrode placement is not restricted with respect to their spatial extent [41,47,108], non-focal stimulation montages might enforce a co-stimulation of various cortical regions [47,109]. In some participants of the present study, the personalization of the tACS montage led to the placement of one set of electrodes over parietal cortex with another set of electrodes of inverted polarity roughly over premotor cortex of the same hemisphere (see Supplement), leading to a co-stimulation of parietal and premotor cortex. Further, previous studies showed that the efficacy of tACS-neuromodulation depends on the intrinsic state of the brain network being involved in the task [90,110,111, see also 112,113]. During covert visuo-spatial attention, the left hemisphere, including the parietal and the premotor cortex, are involved in the modulation of perception and cognition [30,31,87,105,114]. Thus, in line with our results, the same regions might be highly susceptible to subtle neuromodulations, such as low-amplitude tACS.

## Conclusion

In this study, we applied personalized alpha- and gamma-tACS specifically targeting the left and right posterior parietal cortex during covert visuo-spatial attention. We found that left parietal alpha-tACS shifted attention to the left hemifield ipsilateral to electrical stimulation compared to left gamma-tACS. Since no asymmetry was observed for the simulated electric fields between the left and right hemisphere, this lateralization of attention highly supports a tACS-induced modulation of functional properties of the underlying brain networks when targeting the left posterior parietal cortex. Furthermore, ERPs in response to visual stimuli were modulated by alpha versus gamma tACS and were localized in left premotor cortex. These EEG results corroborate the notion of crucial interactions between parietal and premotor cortex during visuo-spatial attention. In addition, a correlation between electric field magnitudes in the left premotor cortex and the behavioral shift of attention indicates that a co-stimulation of the left premotor cortex might contribute to the observed tACS effects. In sum, our results support a role of neuronal alpha activity during covert visuo-spatial attention and suggest that the left dorsal attention network is especially susceptible to subtle tACS-neuromodulations during visuo-spatial attention.

## CRediT authorship contribution statement

**Jan-Ole Radecke:** Conceptualization, Methodology, Software, Investigation, Formal analysis, Writing - original draft, Writing - review & editing, Visualization. **Marina Fiene:** Conceptualization, Methodology, Writing - review & editing. **Jonas Misselhorn:** Conceptualization, Methodology, Writing - review & editing. **Christoph S. Herrmann:** Writing - review & editing, Supervision. **Andreas K. Engel:** Writing - review & editing, Supervision, Resources. **Carsten H. Wolters:** Writing - review & editing, Supervision, Software. **Till R. Schneider:** Conceptualization, Methodology, Software, Writing - review & editing, Project administration, Funding acquisition.

## Acknowledgements

This work was supported by the German Research Foundation (SFB 936 - 178316478 - A3 to TRS and AKE; SPP 1665 - SCHN/1511/1-2 to TRS; SPP1665 - EN 533/13-1 and SFB TRR 169 - B1 to AKE; SPP1665 – WO 1425/5-2 and WO 1425/10-1 to CHW), by the Studienstiftung des deutschen Volkes (to MF) and by the Bundesministerium für Gesundheit (BMG) as project ZMI1-2521FSB006, under the frame of ERA PerMed as project ERAPERMED2020-227 (to CHW). We thank Karin Deazle and Rose Gholami for support with the recruitment of participants and EEG preparation, Jürgen Finsterbusch for technical support and Bettina Schwab for constructive discussions.

