## Supplementary Materials for "Personalized alpha-tACS targeting left posterior parietal cortex modulates visuo-spatial attention and posterior evoked EEG activity"

### Supplementary methods

#### Cued visuo-spatial attention paradigm

During the cued visuo-spatial attention paradigm, participants were presented with one of two sinusoidal auditory cue stimuli (440 Hz or 880 Hz, 100 ms duration, 70 dB SPL) [cf. 1]. Whether the high or the low tone indicated the left or right hemifield was pseudo-randomly assigned across subjects. White-noise bursts (100 ms duration) were presented during the inter-trial interval to indicate incorrect responses. All auditory stimuli were presented by means of bilateral insert earphones (E-A-RTONE 3A, 3M, Indianapolis, IN, USA) and were sampled at 48 kHz. Linear ramps (10 ms) were applied at the beginning and end of auditory stimuli to avoid click perceptions. The amplitude of the 880 Hz tone was scaled to match the perceived loudness of the 440 Hz tone at 70 dB SPL.

Participants were seated in front of an LCD screen (1920 x 1080 pixel, 60 Hz frame rate, 60 cm eye-screen distance) presenting visual stimuli. Each trial was initiated with a red central fixation cross (1° diameter) on a black screen. After a baseline interval (1 to 1.5 s, jittered in steps of 0.2 s) the auditory cue was presented, followed by the cue-stimulus interval (1° diameter red central fixation cross on a black screen; 1 to 1.5 s, jittered in steps of 0.2 s) and the stimulus interval (1 to 1.5 s, jittered in steps of 0.2 s). During the stimulus interval, bilateral random dot kinematograms were presented in the left and right hemifield for maximally 3 s or until a response was indicated via button press. Random dots were presented in round apertures (19° diameter), centered at 13.2° horizontal eccentricity in the left and right hemifield, respectively. White dots were presented on black background (dot size 0.2°, no overlap) at a frame rate of 30 Hz. Dots moved with 11.5°/s. In each frame, a random set of dots was selected according to the coherence level and was moved either up or downwards to provoke a movement illusion.

Individual coherence thresholds were determined using an adaptive procedure [2] defining the coherence at 60% correct responses. For the random dots, two coherence levels were defined at the 60% correct threshold (hard), as well as at the coherence level for the hard condition + 3 % (easy). Coherence thresholds were determined before and after the localizer EEG session. Coherence levels from the two measurements were averaged to determine a robust estimate of the individual coherence levels for the subsequent sessions.

The same coherence levels were used for all four tACS sessions. Across subjects, coherence levels were set at 9.8 ± 4.2 % (hard) and 12.8 ± 4.2 % (easy; M ± SD; Fig. 1B, main manuscript).

Participants indicated upwards or downwards moving random dots via button press with the left and right thumb. Across participants, the mapping of movement direction and left- or right-hand responses was pseudo-randomly assigned.

**FEM headmodel generation**

Tissue conductivities were defined as 0.33 S/m (gray matter), 0.14 S/m (white matter), 1.79 S/m (cerebrospinal fluid), 0.43 S/m (skin), 0.025 S/m (bone spongiosa) and 0.007 S/m (bone compacta), as well as 1.4 S/m for the electrodes [3–5]. Geometry-adapted FEM headmodels were computed using a node shift of 0.33 to ensure that the inferior angles at all element vertices remain convex and the Jacobian determinant in the FEM computations remains positive. Individually registered electrode positions from the EEG layout were aligned to the corresponding headmodel using the fiducial points (nasion and bilateral tragi). Electrode positions were then projected to the closest node of the skin compartment in the FEM headmodel to simulate the electrodes in the framework of a point electrode model [6]. FEM headmodels with *n* nodes (*n* = 3.67 ± 0.31 million, M ± SD; range 3.19 to 4.46 million) were then utilized for EEG source estimation and electric field simulations and computation of personalized tACS montages.

**Behavioral data analysis: *d'* and ln(β)**

In the framework of the signal detection theory, HITs were defined as probabilities of correct UP responses and false positives (FPs) as probabilities of incorrect DOWN responses. Sensitivity index $d'$ was computed as $d' = \mathrm{z(HIT)} - z(\mathrm{FP})$. Response bias ln(β) was computed as $\ln(\beta) = d' * c$ with $\mathrm{c} = 0.5 * [\mathrm{z(HIT)} + \mathrm{z(FP)}]$. The function *z*(*p*), *p* ∈ [0,1], represents the inverse of the cumulative distribution function of the standard normal distribution. To control for possible extreme values, 0.01 was added to HIT and FP equal to 0 and subtracted to HIT and FP equal to 1.

**Definition of stimulation targets**

Continuous EEG data from the EEG localizer were down-sampled to 500 Hz and highpass-filtered at 0.3 Hz half-amplitude cutoff. Data were epoched to cue onset (-0.8 to 1.2 s) and baseline-corrected (-0.6 to 0 s, relative to cue onset). An extended infomax independent component analysis (ICA) [7] was computed after automatically rejecting epochs with improbable events. The weights of ICA components related to eye-blinks, heart-beat and electrical noise were identified by visual inspection of topographies, spectra and temporal dynamics [8] and set to zero (3 ± 2.2 ICs were rejected, M ± SD). Data were re-referenced to common average reference and missing channels were interpolated using a spherical spline. Cross-spectral density was computed for the period from 0.4 to 1 s post cue onset and averaged between 8 to 12 Hz (alpha frequency band).

Using individual FEM leadfields, EEG source reconstructions were computed using exact low-resolution electromagnetic tomography (eLORETA) [9], separately for trials on which participants attended the left or the right hemifield. eLORETA solutions were computed for the EEG activity averaged across the respective time window of interest for each stimulation condition ($\lambda$ = 0.05). Subsequently, a laterality index (LI) was computed as $\frac{attend_{left} - attend_{righ}}{attend_{left} + attend_{right}}$ for every grid point.

Stimulation targets were defined as the summed dipole alpha-activity, defined by the maximal LI in the left hemisphere and the minimal LI in the right hemisphere within an anatomical region of interest along the IPS (inferior and superior parietal cortex; Fig. 2G, amin manuscript) [cf. 10]. Individual T1 images were normalized on the MNI brain to define ROIs. The respective transformation matrix was used to determine anatomical labels for each grid point based on the AAL (Automated anatomical labelling) atlas [11]. Summed dipole positions were estimated as the local maxima (left hemisphere) and minima (right hemisphere) of the LI. Assuming the same neural populations underlying distinct activity patterns during the $attend_{left}$ and $attend_{right}$ conditions, direction vectors were determined as the average direction across the two conditions.

**Individual targeting of tACS montages and electric field simulations**

The Adjoint method [12] implemented in Simbio (SimBio Development Group) was employed for simulation of electric fields. Matrix *A* was computed by combining the individual *m* electrode positions (*m* = 126, Fig. 2H, main manuscript) and the respective six compartment FEM headmodels with *n* nodes (*n* = 3.67 ± 0.31 million, M ± SD; range 3.19 to 4.46 million). Matrix *A* is symmetric to the leadfield in EEG inverse problems due to Helmholtz reciprocity [12,14,15] and can be used to find the optimal weighting of current at the stimulation electrodes $s_{max}$. The linear combination of the weighted current *s* (including the reference electrode) applied to *m* electrodes as a function of *A* determines the current density vector field $j$ at each node $r_n$ for a given stimulation montage:

$$j = As \tag{1}$$

$$\text{with } j = \begin{bmatrix} j(r_1) \\ j(r_2) \\ \vdots \\ j(r_n) \end{bmatrix}, A = \begin{bmatrix} a_1(r_1) & a_2(r_1) & \dots & a_m(r_1) \\ a_1(r_2) & a_2(r_2) & \dots & a_m(r_2) \\ \vdots & \vdots & \ddots & \vdots \\ a_1(r_n) & a_2(r_n) & \dots & a_m(r_n) \end{bmatrix} \text{ and } s = \begin{bmatrix} s_1 \\ s_2 \\ \vdots \\ s_{m-1} \\ -\sum s_m \end{bmatrix}$$

The Distributed Constrained Maximum Intensity (DCMI) is expressed as:

$$s_{max} = \arg\max_s < Cs, o_t > -\lambda \|s\|_2 \tag{2}$$

subject to $\|s\|_1 \leq 2i_{Total}$ and $\|s\|_\infty \leq i_{Limit}$

with $C = [a_1(r_t), a_2(r_t), \dots, a_m(r_t)]$

where $r_t$ denotes the index of the target node and $o_t$the target vector orientation.

C is the submatrix of A that reflects the mapping of the electrode currents to the current density at the target vector. We chose $i_{Total}$ = 2 mA to fulfill the safety constraint and $i_{Limit}$ = 0.95 mA to reduce the current applied to each electrode and thereby reduce potential tactile perception of the stimulation. Regularization was set to $\lambda$ = 250 to slightly distribute the injected currents across stimulation electrodes. The number of electrodes was fixed to six electrodes (three electrodes of each polarity) in a two-step procedure. First, DCMI optimization was computed including all possible electrode positions from the cap layout. Next, three electrodes of each polarity with the maximum weights were selected for a second computation of the DCMI, resulting in the final tACS montage. Beside the high directionality, DCMI offers potentially reduced side effects and skin sensations, the DCMI solution is unique due to the fact that the regularization adds convexity to the optimization cost function, and there are no jumps of injection currents between electrodes due to for example tiny changes in volume conduction modeling or sensor registration [16–19].

EEG cap positioning was related to the nasion and inion, as well as the left and right tragus to ensure a correct transfer between the digitally optimized electrode positions and the application of personalized tACS. During every session, electrode positions were registered to validate cap positioning.

**EEG pre-processing**

If not indicated otherwise, Kaiser-windowed zero-phase FIR filter were applied with a maximum passband deviation of 0.001 (kaiser-β = 5.65) and a transition bandwidth of 2 Hz. Half-amplitude cutoff (Hz) is reported and separate low- and highpass filters were applied for bandpass-filtering [20].

ICA was used to reject stereotypical artifacts from the EEG data. An extended infomax ICA [7] was computed to address low-frequency-specific artifacts. For the ICA-preprocessing, the unepoched data were filtered between 1-35 Hz. Dummy epochs of 1s were created and artifactual channels and epochs were rejected semi-automatically (joint probability and kurtosis criteria, 5 SD; raw data inspection), before running the ICA. The computed ICA weights were then applied to the data that were epoched to the cue and stimulus onset, respectively.

**EEG source estimation**

eLORETA was utilized to estimate sources of EEG sensor data [9]. eLORETA estimates a spatial filter matrix $W_S$ to solve the equation:

$$j(t) = W_S V(t) \tag{3}$$

where *j(t)* represents the dipole moment, and *V(t)* the vector sensor activity as a function of time, respectively. With eLORETA aiming to minimize the product $H = W_S L^T$, the spatial filter is computed as follows:

$$W_S = [L^T (L C^{-1} L^T + \lambda H)^{+} L]^{1/2} \tag{4}$$

where *L* represents the leadfield, *C* is the data covariance matrix, $\lambda$ is the regularization parameter, superscript + denotes the Moore-Penrose pseudo-inverse and *H* is the centering matrix of the surface Laplacian. Single orientation solutions were computed using singular value decomposition [21].

**Parameter computation for electric field simulations**

Three parameters were computed, describing the individual electric field simulations with respect to the stimulation target. First, the parallelity ($E_{par}$) between the stimulation target orientation vector $u_t$ and the target electric field orientation vector $v_t$ was defined as the absolute of the dot product of the two vectors $E_{par} = |u_t \cdot v_t|$. This parameter results in 1 if the electric field and the target are perfectly parallel or antiparallel within the target, and in 0 if the orientations of the electric field and the target vectors are orthogonal. Second, target intensity ($E_{target}$) was defined as the target electric field vector length (GRAY matter only), corrected for the parallelity with the stimulation target vector: $E_{target} = |e_t|\, E_{par}$, with $|e_t| = \sqrt{e_{t\,1}^2 + e_{t\,2}^2 + e_{t\,3}^2}$ (directionality [16]). Third, to estimate the spatial extent of the electric field relative to the stimulation target ($E_{extent}$), the 0.95-percentile electric field magnitude (GRAY and WHITE matter) was computed as a function of the Euclidean distance to the stimulation target (5 to 100 mm radius, in steps of 5 mm), normalized to the maximum value. Spatial extent was quantified as the distance (in mm) at 50% area under the curve of this function [3]. For illustration, individual electric fields were interpolated on a common MNI cortical grid and averaged across subjects for $IPS_L$ and $IPS_R$, respectively. Personalized stimulation montages and the induced electric fields are presented in addition to the average and standard deviation of the electric field magnitudes (Suppl. Fig. 1).

### Supplementary results

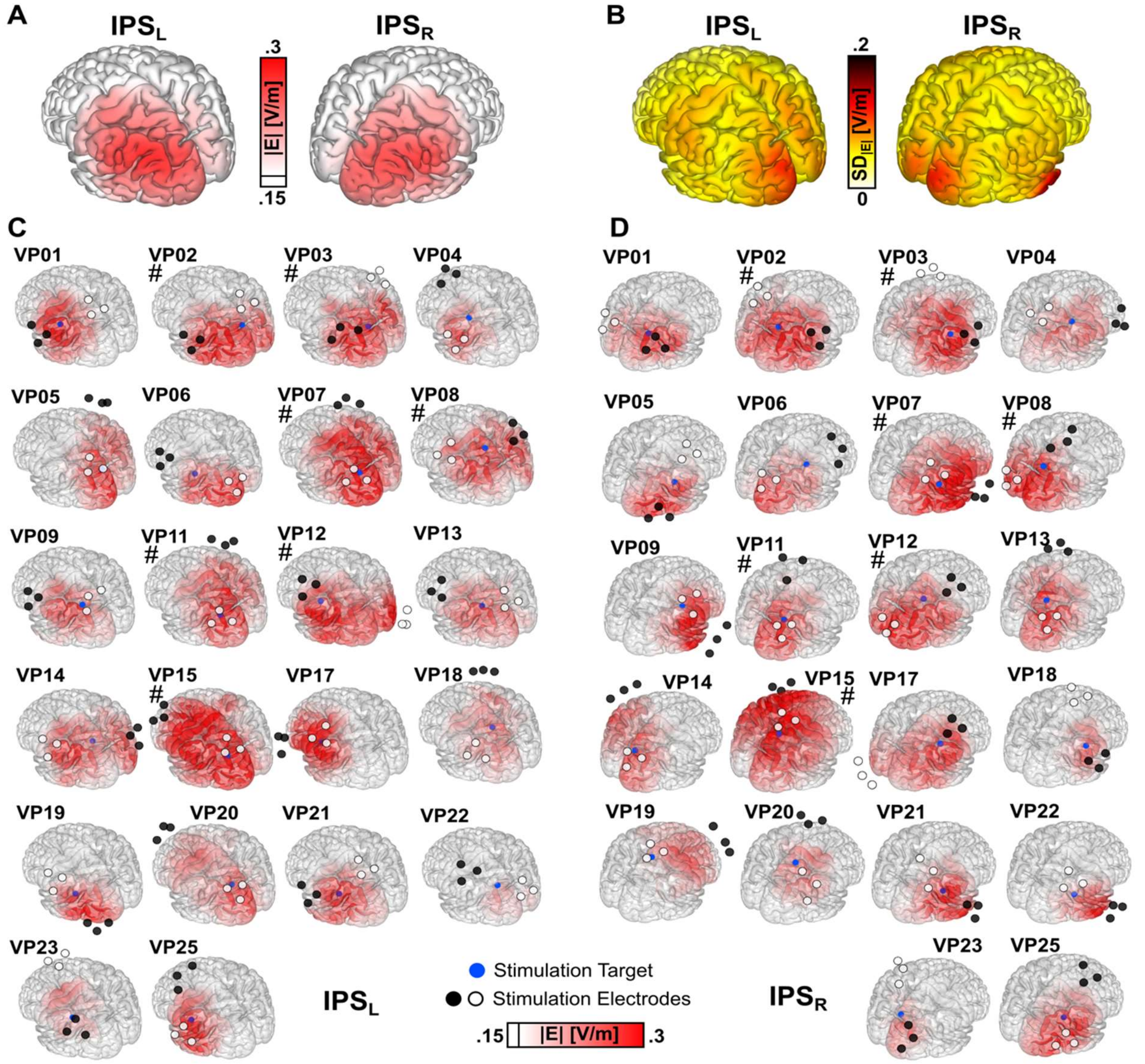


**Supplementary figure 1. Single-subject electric field simulations. A)** Average electric field magnitudes for tACS montages targeting the left ($IPS_L$) and right ($IPS_R$) parietal cortex, thresholded at 0.15 V/m. **B)** Standard deviation of electric fields across participants for $IPS_L$ and $IPS_R$. **C)** Individual tACS-montages, stimulation target locations and electric field magnitudes interpolated on the cortical surface of the MNI brain for tACS montages targeting $IPS_L$ (left) and $IPS_R$ (right), respectively. Seven participants received tACS at 2 mA (marked by #), all other participants received 1.5 mA. White and black circles represent electrodes with the same polarity. The exact currents applied at each electrode were neglected here for illustrative reasons (note: maximum current per electrode was 1 mA).

#### Analysis of electric field simulations

The illustration in the main manuscript (Fig. 2G, main manuscript) indicates the validity of the approach, in which a relatively extended brain activity can be estimated by single target vectors (representing the center of the activation of interest), that in turn lead to an extended electric field when applying the optimization of tACS montages. Personalized electrode montages across all participants are centered in posterior scalp regions along the IPS, with electrodes placed at left or right scalp regions for targeting $IPS_L$ and $IPS_R$, respectively (Fig. 2I, main manuscript). The electrode polarities (please note that anodes and cathodes change polarity over time during tACS application) are generally oriented along the anterior-posterior axis, indicating a rather tangential orientation of the stimulation targets [3]. Only few electrodes were placed in remote frontal or occipital scalp regions. Presumably, these electrodes controlled the targeted electric fields in case of more radial or deeper stimulation targets, that would require more distant stimulation montages when using algorithmic optimization to increase target intensity [3,5].

**Supplementary table 1.** Electric field intensities across different tissue types. The average electric fields intensity across the maximum 10.000 nodes are reported for each tissue type (M ± SD) and for electric fields simulated for targeting $IPS_L$ and $IPS_R$, respectively.

| E-Field | *$IPS_L$ [V/m]* | *$IPS_R$ [V/m]* |
|---|---|---|
| **SKIN** | 8.62 ± 1.28 | 8.61 ± 1.33 |
| **BONE** | 15.27 ± 2.83 | 14.86 ± 2.5 |
| **CSF** | 0.2 ± 0.04 | 0.21 ± 0.04 |
| **GRAY** | 0.36 ± 0.06 | 0.37 ± 0.08 |
| **WHITE** | 0.4 ± 0.08 | 0.4 ± 0.07 |

Electric field magnitudes were different across tissue types. Descriptive electric field magnitudes ($E_{kmax}$) across tissue types and stimulation targets ($IPS_L$, $IPS_R$, as well as the follow-up of the main effect of Tissue are shown in Supplementary Table 1 and 2, respectively.

**Supplementary table 2.** Paired t-tests across different tissue types, based on the average electric field intensity across the maximum 10.000 nodes. Electric field intensities were averaged across conditions targeting $IPS_L$ and $IPS_R$. *t*-values, degrees of freedom, corrected *p*-values and Cohen' *d* are reported. Asterisks indicate significant differences.

| Tissue | $t_{df}$ | *df* | *p* | *Cohen's d* |
|---|---|---|---|---|
| **SKIN vs BONE** | -17.56 | 21 | ***< .001 **** | 3.75 |
| **SKIN vs CSF** | 31.49 | 21 | ***< .001 **** | 6.71 |
| **SKIN vs GRAY** | 31.15 | 21 | ***< .001 **** | 6.64 |
| **SKIN vs WHITE** | 31.5 | 21 | ***< .001 **** | 6.72 |
| **BONE vs CSF** | 27.24 | 21 | ***< .001 **** | 5.81 |
| **BONE vs GRAY** | 27.1 | 21 | ***< .001 **** | 5.77 |
| **BONE vs WHITE** | 27.22 | 21 | ***< .001 **** | 5.8 |
| **CSF vs GRAY** | -21.18 | 21 | ***< .001 **** | 4.5 |
| **CSF vs WHITE** | -22.65 | 21 | ***< .001 **** | 4.83 |
| **GRAY vs WHITE** | -4.68 | 21 | .129 | 1 |

### Cue-related alpha power lateralization was not modulated by tACS

Sensor level time-frequency dynamics were determined using an FFT approach for -1 to 1 s relative to the cue and stimulus onset, respectively. For the low-frequency data, total power was determined for frequencies between 4 to 25 Hz in steps of 1 Hz. Total power was computed using sliding time-windows in steps of 50 ms, multiplied by a single Hanning-taper. Window length was adapted between 500 to 250 ms to capture two cycles for each frequency below 8 Hz (frequency smoothing: ± 1 to 2 Hz). For frequencies between 8 to 25 Hz, window length was fixed to 250 ms (frequency smoothing: ± 2 Hz). Results were averaged across

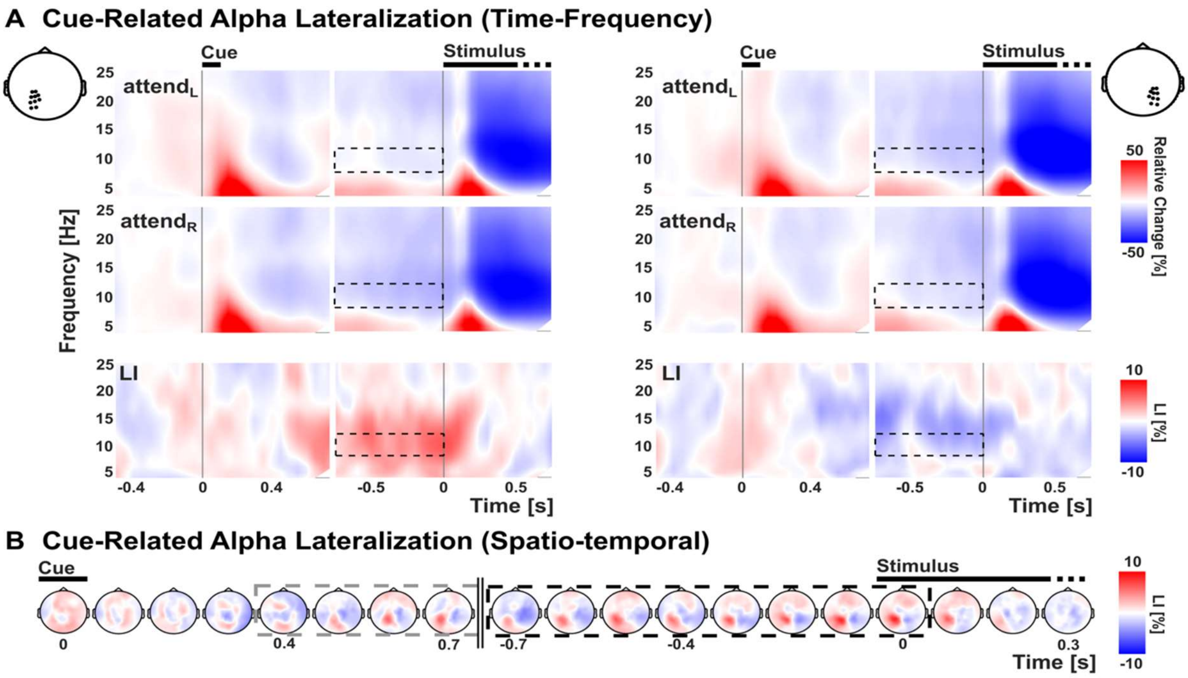


**Supplementary figure 2. Cue-related alpha power lateralization during the localizer. A)** Time-frequency representation of low-frequency EEG data averaged across left and right posterior electrode clusters. Time-frequency data relative to cue and stimulus-onset for attend$_L$, attend$_R$, and the laterality index (LI). The time-frequency window for alpha power is marked by black dashed boxes (8-12 Hz, -0.75 to 0 s relative to stimulus onset). **B)** Spatio-temporal representation of alpha power lateralization relative to cue and stimulus onset. Grey dashed lines represent the time-window used for localizing parietal alpha power. Black dashed lines represent the time-window for analysis of alpha lateralization. Both time windows are partly overlapping, due to the jittered cue-stimulus interval and represent the ongoing lateralization of alpha power in posterior sensors during the cue-stimulus interval.

electrodes for two electrode clusters of interest in sensor space. Time-frequency dynamics were computed relative to baseline for each attention side as $\frac{attend_{left} - baseline}{baseline}$ and $\frac{attend_{right} - baseline}{baseline}$. LI was computed as $\frac{attend_{left} - attend_{right}}{attend_{left} + attend_{right}}$ for every grid point (Suppl. Fig. 2). Due to the hypotheses concerning the occurrence and tACS-modulation of alpha activity during the cued visuo-spatial attention paradigm, statistical analysis was conducted in pre-defined time-frequency windows (8-12 Hz, -0.75 to 0 seconds, relative to stimulus onset).

We did not observe a tACS-modulation of alpha power, neither on sensor-level (main manuscript), nor on source-level in the superior occipital cortex (not shown), the targeted parietal cortex (Fig. 3, main manuscript), the middle occipital cortex (Suppl. Fig. 3A), the area typically showing the strongest alpha-lateralization during visuo-spatial attention [22], or the superior occipital cortex (Suppl. Fig.3B).

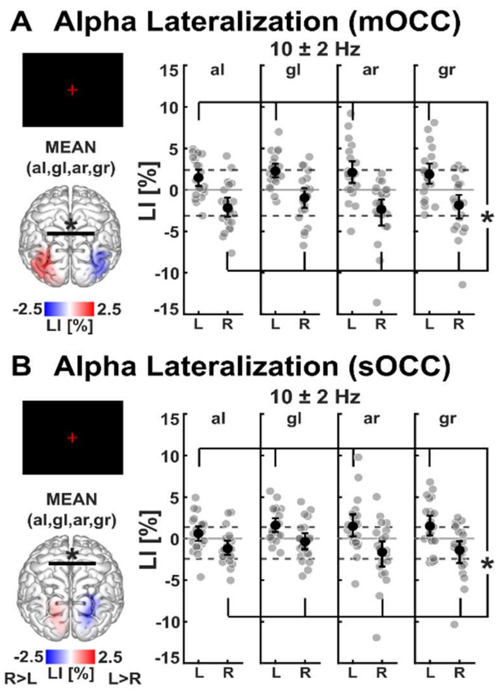


**Supplementary figure 3. No tACS modulation of alpha power lateralization in EEG source estimates.** In addition to the analysis of alpha power in the parietal cortex (see main manuscript), occipital source regions of interest were assessed for tACS-modulations of cue-related alpha power lateralization. **A)** Analysis of alpha total power lateralization in middle occipital cortex (mOCC), and **B)** superior occipital cortex (sOCC) did not reveal tACS modulation effects. However, across all four tACS sessions, significant differences between attend$_L$ and attend$_R$ were observed in both regions of interest. * represent p < 0.05. Individual values and bootstrapped mean ± 95%-confidence interval are depicted, respectively. Dashed lines represent the descriptive mean LI-values that were observed during the localizer task.

**Supplementary table 4.** Wilcoxon signed-rank test results comparing alpha source power between attend$_L$ and attend$_R$ during the localizer. Test results are reported for regions of interest in left and right intraparietal sulcus (IPS), middle occipital cortex (mOCC) and superior occipital cortex (sOCC). Subscript letters indicate the left (L) and right (R) hemisphere, respectively. Corrected p-values are reported. Asterisks indicate significant differences. Median power values and interquartile ranges [25th to 75th percentile] are reported.

| Localizer | $z_N$ | N | *p* | *\|r\|* | *Median Power att$_L$* | *Median Power att$_R$* |
|---|---|---|---|---|---|---|
| **IPS$_L$** | -1.70 | 22 | .088 | 0.36 | 137 [83 to 334] | 129 [86 to 302] |
| **IPS$_R$** | -2.97 | 22 | ***.015 **** | 0.63 | 119 [58 to 300] | 147 [61 to 310] |
| **sOCC$_L$** | -2.06 | 22 | .079 | 0.44 | 132 [67 to 724] | 135 [71 to 580] |
| **sOCC$_R$** | -3.07 | 22 | ***.013 **** | 0.65 | 136 [90 to 681] | 167 [102 to 764] |
| **mOCC$_L$** | -2.65 | 22 | ***.024 **** | 0.56 | 162 [81 to 521] | 148 [73 to 403] |
| **mOCC$_R$** | -2.84 | 22 | ***.018 **** | 0.61 | 148 [99 to 632] | 208 [107 to 643] |

The statistical analysis of the source-level alpha power lateralization during the localizer revealed an interaction effect of ROI * Hemisphere * Attention Side interaction ($F_{1.5,32.1}$ = 6.73, $p$ = .007, $\eta_p^2$ = .243). No tACS-modulation of source-level alpha power lateralization was observed. However, the ROI * Hemisphere * Attention Side interaction was confirmed across all stimulation conditions ($F_{1.2,25.5}$ = 4.15, $p$ = .045, $\eta_p^2$ = .165). Post-hoc test results (Wilcoxon signed-rank) are depicted in supplementary table 4 (localizer) supplementary table 5 (tACS), indicating a significant lateralization of alpha power.

**Supplementary table 5.** Wilcoxon signed-rank test results comparing alpha source power between $attend_L$ and $attend_R$ for the tACS sessions (averaged across the four stimulation conditions). Test results are reported for regions of interest in left and right intraparietal sulcus (IPS), middle occipital cortex (mOCC) and superior occipital cortex (sOCC). Subscript letters indicate the left (L) and right (R) hemisphere, respectively. Corrected p-values are reported. Asterisks indicate significant differences. Median power values and interquartile ranges [25th to 75th percentile] are reported.

| ROI | $z_N$ | N | *p* | *\|r\|* | *Median Power $att_L$ [µV²]* | *Median Power $att_R$ [µV²]* |
|---|---|---|---|---|---|---|
| **$IPS_L$** | -2.91 | 22 | ***.007 **** | 0.62 | 204 [75 to 513] | 203 [75 to 481] |
| **$IPS_R$** | -3.59 | 22 | ***.002 **** | 0.76 | 158 [73 to 421] | 186 [71 to 451] |
| **$sOCC_L$** | -2.39 | 22 | ***.017 **** | 0.51 | 231 [77 to 816] | 215 [76 to 804] |
| **$sOCC_R$** | -3.07 | 22 | ***.007 **** | 0.65 | 218 [117 to 889] | 237 [118 to 941] |
| **$mOCC_L$** | -3.43 | 22 | ***.003 **** | 0.73 | 249 [107 to 617] | 216 [103 to 585] |
| **$mOCC_R$** | -3.23 | 22 | ***.005 **** | 0.69 | 217 [148 to 594] | 236 [150 to 673] |

### Stimulus-related gamma power response was not modulated by tACS

The EEG dataset was split into a low frequency dataset (0.3 - 35 Hz), by the application of a lowpass-filter at 35 Hz (reported in the main manuscript), and a high-frequency dataset (16 - 250 Hz), by the application of a highpass-filter at 16 Hz [cf. 23]. In addition to the horizontal and vertical EOG described in the main manuscript, radial EOG (EEG/EOG channels around the eyes referenced against centro-parietal channels) was computed in the high-frequency dataset to determine the occurrence of saccadic spike potentials [24]. In parallel to the ICA that was applied to the low-frequency dataset, an additional extended infomax ICA [7] was computed to address high-frequency-specific artifacts. For the ICA-preprocessing, the unepoched data were filtered between 16-250 Hz and dummy epochs of 1 s were created. Dummy epochs of 1 s were created and artifactual channels and epochs were rejected automatically (joint probability and kurtosis criteria, 5 SD; raw data inspection), before running the ICA. The computed ICA weights were then applied to the data that were epoched to the cue and stimulus onset, respectively. For the high-frequency dataset, components related to 50 Hz line noise, saccadic spike potentials and electrical muscle artifacts were identified based on topographies, spectra and temporal dynamics, as well as on the relation of each component to the radial EOG signal [23–26] and the respective ICA weights were set to zero ($54.6 \pm 5.2$ ICs were rejected). Finally, the data were re-referenced to common average reference and missing channels were interpolated using a spherical spline.

For the high-frequency data, total power as a function of time was determined for ten center frequencies with 1/4 octave step-size between 26 to 128 Hz using a sliding window of 250 ms (in steps of 50 ms) and orthogonal Slepian tapers to adapt frequency smoothing with increasing frequency (3 to 16 tapers, frequency smoothing: $\pm$ 8 to 34 Hz). Results were averaged across electrodes for two posterior electrode clusters of interest in sensor space. Time-frequency dynamics were computed relative to baseline for each attention side as $\frac{attend_{left} - baseline}{baseline}$ and $\frac{attend_{right} - baseline}{baseline}$. LI was computed as $\frac{attend_{left} - attend_{right}}{attend_{left} + attend_{right}}$ for every grid point. These time-frequency representations of gamma power are shown in Suppl. Fig. 4A.

Stimulus-related gamma-band power was determined after stimulus onset (0.2 to 0.6 s relative to stimulus onset). Gamma-power was computed for a center frequency of 70 $\pm$ 19 Hz using 14 Slepian tapers [cf. 22]. Results were averaged across electrodes for two posterior electrode clusters of interest in sensor space. eLORETA was utilized to estimate stimulus-related gamma total power (0.2 to 0.6 s relative to stimulus onset) along the orientation of largest power using singular value decomposition [21]. LI was computed as $\frac{attend_{left} - attend_{right}}{attend_{left} + attend_{rig}}$ for every grid point. A repeated-measures ANOVA was computed to test tACS-modulation of sensor gamma total power including the factors Stimulation Frequency [alpha, gamma], Stimulation Side [$IPS_L$, $IPS_R$], Electrode Cluster [lp, rp], and Attention Side [$attend_L$, $attend_R$]. Sensor-level analysis of stimulus-related gamma total power during the localizer (Suppl. Fig. 4A) revealed a significant main effect of Electrode Cluster ($F_{1,21}$ = 9.57, $p$ = .006, $\eta_p^2$ = .313), indicating higher gamma power in the left, compared to the right posterior electrode cluster, irrespective of attention side. No main or interaction effect including Attention Side was observed (all $p$ > .385). Sensor-level analysis of stimulus-related gamma total power during the four tACS sessions (Suppl. Fig. 4B) revealed a significant Stimulation Frequency * Stimulation Side * Electrode Cluster interaction ($F_{1,21}$ = 5.41, p = .03, $\eta_p^2$ = .205; all other main or interaction effects: p > .104). However, follow-up paired t-tests did not confirm a significant difference between electrode clusters, specific to stimulation condition (averaged across attention conditions; all p > .9). Since no tACS-modulation of gamma power and no stimulus-related lateralization was revealed by this analysis, gamma power was not further assessed on source-level.

Relative to baseline, a descriptive gamma-response was observed (Suppl. Fig. 4A) that can be related to stimulus-related neural processing of the random dots [10]. This response was not further analyzed in the current study.

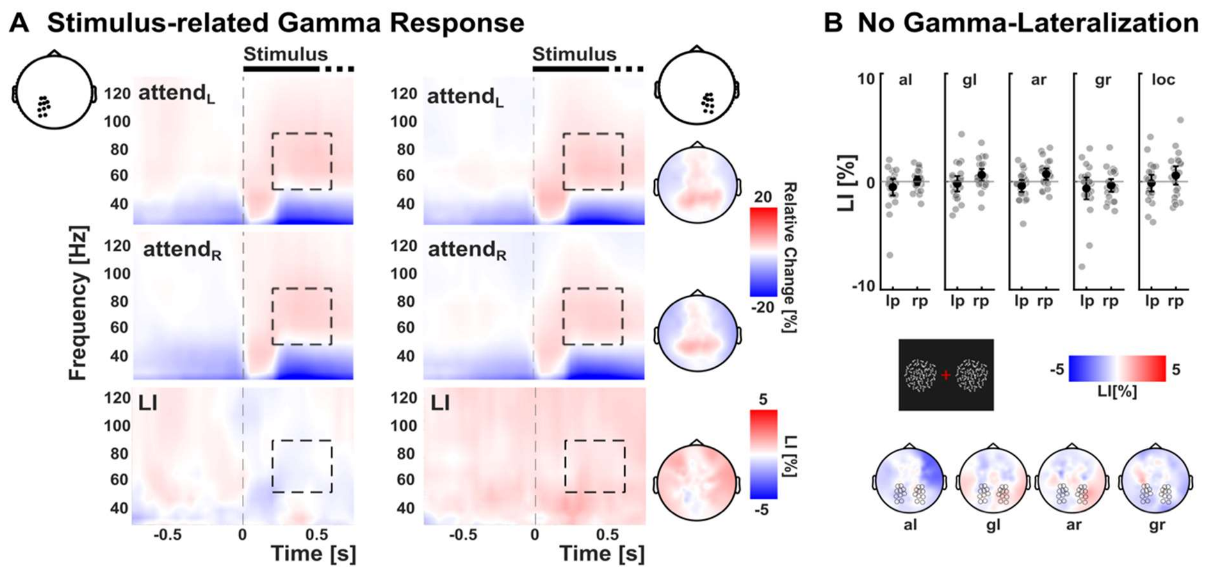


**Supplementary figure 4. Stimulus-related gamma activity. A)** Time-frequency representation of ongoing stimulus-related high-frequency EEG activity in a left and right posterior electrode cluster for attend$_L$, attend$_R$ and the laterality index (LI) during the localizer task. An increase in posterior gamma activity relative to baseline can be observed in between 70 ± 19 Hz (0.2 to 0.6 s) for both attend$_L$ and attend$_R$. Topographical representations of the same time-frequency window are shown on the right, putatively reflecting the induced gamma activity in response to the bilateral random dot kinematograms. No lateralization of ongoing gamma activity (bottom), and **B)** no tACS-modulation of gamma activity was observed on sensor-level.

### Cue ERP analysis and descriptive stimulus ERPs

In addition to visual stimulus ERPs, auditory cue ERPs were computed in response to cue stimuli (-0.2 to 0.75 s, relative to cue onset), separately for attending to the left and right hemifield for the localizer and all four tACS sessions. Epochs were averaged and baseline-corrected (-0.2 to 0 s). Difference ERPs were computed by subtracting ERPs of attend$_R$ from ERPs of attend$_L$ (attend$_L$ - attend$_R$). Non-parametric cluster permutation test [27] for the localizer cue ERPs was applied for the time-window 0 to 0.75 s relative to cue-onset and all 126 EEG sensors (paired *t*-tests, 1000 permutations, $\alpha_{cluster}$ = 0.05, $\alpha$ = 0.05, two-sided).

Descriptive ERP components were defined based on the localizer grand average ERP (averaging attend$_L$ and attend$_R$). Mean amplitudes as well as 50%-area latencies were computed. Auditory cue ERP components were defined in a central electrode cluster in the latency range from 0.104 to 0.144 s (N1) and from 0.186 to 0.246 s (P2). Visual stimulus ERP components were defined in a left and right posterior electrode cluster in the latency range from 0.1 to 0.16 s (P1), 0.186 to 0.246 s (N1), 0.272 to 0.312 s (P2), and from 0.352 to 0.552 s (P3).

**Supplementary table 4.** Descriptive amplitudes and latencies of observed ERP components. Mean amplitudes and 50%-signed area latencies are reported (M ± SD).

| ERP Components | *Amplitudes [µV]* | *Latencies [ms]* |
|---|---|---|
| **Auditory N1** | -1.88 ± 1.23 | 123 ± 6 |
| **Auditory P2** | 3.19 ± 1.5 | 201 ± 4 |
| **Visual P1** | 0.92 ± 0.95 | 120 ± 10 |
| **Visual N1** | -0.53 ± 1.77 | 207 ± 13 |
| **Visual P2** | 0.17 ± 1.89 | 280 ± 7 |
| **Visual P3** | 0.49 ± 0.98 | 392 ± 26 |

Cue ERPs showed clear evoked activity in response to the auditory cues. Auditory N1 and P2 components were observed with fronto-centrally pronounced topographies (Suppl. Fig. 5, Suppl. Tab. 4). During the localizer, we observed a significant positive effect ($p$ = .005) based on a cluster of parieto-central electrodes ranging from 284 to 512 ms ($n_{clustersize}$ = 1595), indicating increased auditory ERP amplitudes for $attend_L$, compared to $attend_R$ (Suppl. Fig. 5B). Similar cue-related lateralization of slow evoked responses during visuo-spatial tasks [28,29] have been linked to the lateralization of oscillatory alpha activity typically observed in visuo-spatial attention paradigms [30]. However, for the same spatiotemporal cluster, no main or interaction effects were observed for auditory cue ERP amplitudes during the four tACS-sessions (repeated-measures ANOVA; all $p$ > .104).

Stimulus-related ERPs reflect activity in response to the random dots. On a descriptive level, visual ERP amplitudes during the localizer were strongly varying between attention conditions ($attend_L$, $attend_R$) and electrode clusters (lp, rp), as illustrated in Suppl. Fig. 5C. During the tACS-sessions, a similar pattern of stimulus-related slow amplitude variations occurred. These amplitude variations were reduced during the left alpha-tACS condition Suppl. Fig. 5D, indicating a specific tACS-modulation of stimulus-related ERPs as described in the main manuscript. Descriptively, both an increase in amplitudes of ERPs to attended stimuli in the right (ipsilateral) hemifield and reduced amplitudes to attended stimuli in the left (contralateral) hemifield seems to drive the reduced difference ERP induced by left alpha-tACS (Suppl. Fig. 5D).

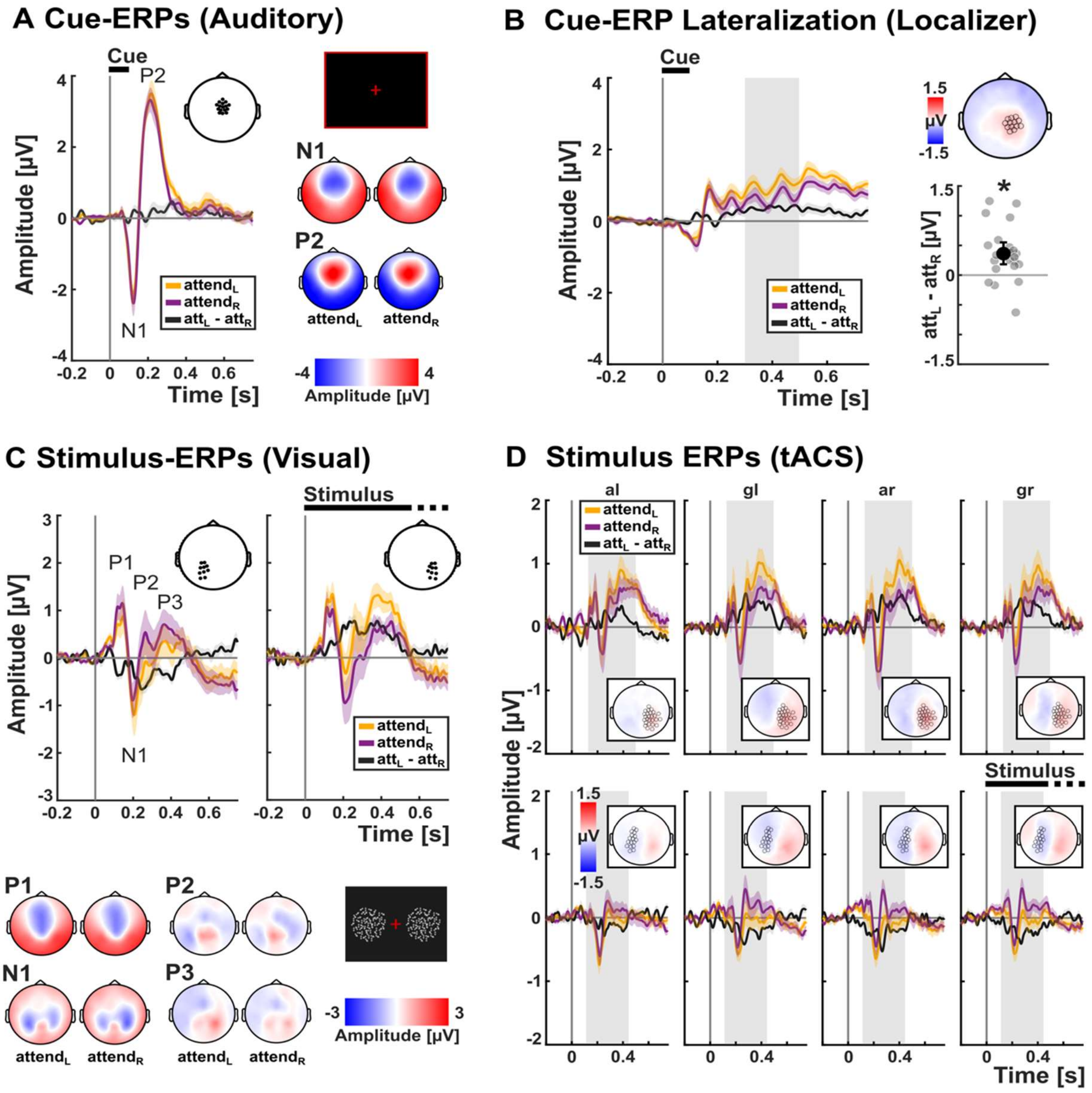


**Supplementary figure 5. Cue-related and stimulus-related ERPs. A)** Localizer: Cue-related auditory ERP time-series (left) and topographies (right) show clear N1 and P2 components in a central electrode cluster that were similar between attend$_L$ and attend$_R$. **B)** Localizer: A significant attention effect of cue-related amplitudes between attend$_L$ and attend$_R$ was identified in a right central electrode cluster. Time-series, topographical representation and mean amplitudes (individual values (attend$_L$-attend$_R$) and bootstrapped mean ± 95%-confidence interval) of the cluster are presented. No cue-related tACS modulation of ERP amplitudes was observed for this cluster (not shown). **C)** Localizer: Stimulus-related visual ERP time-series in a left and right posterior electrode cluster (top) and topographies (bottom) showing a clear amplitude lateralization between attend$_L$ and attend$_R$ across the latency range of P1, N1, P2 and P3 ERP components. **D)** tACS: Stimulus-related visual ERP time-series and topographies in the right posterior (top) and left central-posterior electrode clusters (bottom) during all four tACS sessions. Spatio-temporal clusters are identical to those identified during the localizer experiment.
